# Multi-omic human neural organoid cell atlas of the posterior brain

**DOI:** 10.1101/2025.03.20.644368

**Authors:** Nadezhda Azbukina, Zhisong He, Hsiu-Chuan Lin, Malgorzata Santel, Bijan Kashanian, Ashley Maynard, Tivadar Török, Ryoko Okamoto, Marina Nikolova, Sabina Kanton, Valentin Brösamle, Rene Holtackers, J. Gray Camp, Barbara Treutlein

**Affiliations:** Department of Biosystems Science and Engineering, ETH Zürich, Basel, Switzerland; Max Planck Institute for Evolutionary Anthropology, Leipzig, Germany; Institute of Human Biology (IHB), Roche Pharma Research and Early Development, Roche Innovation Center Basel, Basel, Switzerland; Biozentrum, University of Basel, Basel, Switzerland

## Abstract

Patterning of the neural tube establishes midbrain and hindbrain structures that coordinate motor movement, process sensory input, and integrate cognitive functions. Cellular impairment within these structures underlie diverse neurological disorders, and *in vitro* organoid models promise inroads to understand development, model disease, and assess therapeutics. Here, we use paired single-cell transcriptome and accessible chromatin sequencing to map cell composition and regulatory mechanisms in organoid models of midbrain and hindbrain. We find that existing midbrain organoid protocols generate ventral and dorsal cell types, and cover regions including floor plate, dorsal and ventral midbrain, as well as adjacent hindbrain regions, such as cerebellum. Gene regulatory network (GRN) inference and transcription factor perturbation resolve mechanisms underlying neuronal differentiation. A single-cell multiplexed patterning screen identifies morphogen concentration and combinations that expand existing organoid models, including conditions that generate medulla glycinergic neurons and cerebellum glutamatergic subtypes. Differential abundance of cell states across screen conditions enables differentiation trajectory reconstruction from region-specific progenitors towards diverse neuron types of mid- and hindbrain, which reveals morphogen-regulon regulatory relationships underlying neuronal fate specification. Altogether, we present a single-cell multi-omic atlas and morphogen screen of human neural organoid models of the posterior brain, advancing our understanding of the co-developmental dynamics of regions within the developing human brain.

## Introduction

The mesencephalon (midbrain) and rhombencephalon (hindbrain) are two of the three primary brain vesicles emerging from the anterior part of the neural tube. These regions develop into the cerebral peduncles, tegmentum and tectum (midbrain) and medulla, pons, and cerebellum (hindbrain) of the human brain. Mid- and hindbrain function to coordinate motor movements and process sensory inputs^1^, and their impairment is associated with diverse disorders^2^ including malformations and Parkinson’s disease. However, these brain regions remain understudied, with restricted focus on developing a multicellular model that recapitulates their cell type diversity.

Human brain organoids are three-dimensional developmental brain models generated from human embryonic stem (ES) cells or induced pluripotent stem (iPS) cells. They have served as an accessible gateway to obtain diverse human neural system cell phenotypes and recapitulate intercellular connections and interactions^3^. A variety of protocols have been established to direct differentiation into specific and broad brain regions, serving as functional models to understand mechanisms that underlie human brain development^4,5^ and neurodevelopmental disorders^6,7^. So far, numerous protocols have been reported to generate midbrain^8–14^, hindbrain^15,16^, cerebellum^17,18^, and brainstem fate organoids^19,20^. However, the cell type composition and molecular feature profiling of most of those protocols remains very limited. Recent efforts on integration of existing single-cell RNA-sequencing (scRNA-seq) data of different neural organoid protocols and comparison to primary human brain atlases have also revealed the under-representation of posterior brain region neuron populations in organoids^21^. Furthermore, we lack a full understanding of the molecular mechanisms underlying posterior brain cell type specification and development.

Morphogens are secreted signaling molecules which direct the development of cell types and tissues. They have been widely used during neural organoid cultures to guide their development towards certain brain regions. Most of the morphogen cocktails to guide organoid developments are derived from previous studies on non-human animal models. Recently, systematic explorations of morphogen-induced patterning of human neural organoids emerged as a new strategy for the targeted development and optimization of organoid models^22–24^. By examining the effects of modulations of SHH, BMPs, FGFs, WNTs, and retinoid acid signaling pathways, these studies have identified the core morphogen-modulated effects on the axial and regional specification of human neural organoids. However, conditions explored by the existing studies were biased to generate forebrain cell types, or lack of well-defined conditions. It remains unclear how these signalling molecules influence cell type diversity generation of the posterior brain and to what extent we could guide their specification.

In this study, we generated a single-nucleus multi-omic atlas profiling transcriptome and accessible chromatin from the same single cells over a time course of posterior brain organoid development. A gene-regulatory network was inferred to determine cell type-specific transcription factor regulons, which were perturbed using single-cell CRISPR perturbation experiments to understand their requirement for cell state and regional fate establishment. To derive organoids with novel brain regional identities, we conducted a multiplexed morphogen patterning screen with single-cell transcriptomic readout of individual organoids in triplicates, testing 48 concentrations and combinations of 10 morphogens involved in brain patterning. We identified conditions driving the emergence of under-represented cell types, such as medulla glycinergic neurons and cerebellum glutamatergic neurons, thereby expanding existing mesencephalon and rhombencephalon organoid models. Together, our findings present the first comprehensive single-cell multi-omic atlas and morphogen screen of human posterior brain organoid development, advancing our understanding of its complexity and developmental dynamics.

## Results

### Multi-omic cell atlas of posterior brain organoids identifies features recapitulating primary brain development

We generated a time course single-nucleus multi-omic dataset with paired measurement of transcriptomes and accessible chromatin to map cellular composition and gene regulatory dynamics in two previously established protocols modeling midbrain development^13,14,25^ (Fig. 1, Extended Data Fig. 1, Supplementary Table 1). These protocols use the WNT pathway activator CHIR-99021 (CHIR), sonic hedgehog (SHH), and fibroblast growth factor 8 (FGF-8) for patterning of neural epithelium in a continuous (protocol 1) or sequential (protocol 2) manner, respectively (Fig. 1a-c). The dataset incorporates 104,452 cells collected from 5 time points between day 7 and day 120 of organoid development from 3 human induced pluripotent stem cell (iPSC) lines, covering periods of regionalization, neurogenesis, and maturation. We observed time-dependent trajectories from early neuroepithelial cells via neural progenitor cells (NPCs) to multiple neuronal cell populations including dopaminergic, glutamatergic, GABAergic and glycinergic neurons (Fig. 1d-g; Extended Data Fig. 2b-d). Interestingly, both protocols generated a substantial proportion (57%) of neurons with hindbrain rather than midbrain signatures including glycinergic neurons and Purkinje cells, suggesting successful posteriorization without restriction to midbrain cell types. Notably, the two protocols generated different neuronal populations (Fig. 1e-g). Protocol 1 produced the majority (89%) of dopaminergic neurons with multiple subtypes, including *KCNJ6*+ A9-like dopaminergic neurons and *SOX6*+ *OTX2*+ populations (Extended Data Fig. 2e-g). Protocol 2 produced the majority (82%) of glutamatergic neurons as well as of midbrain GABAergic neurons (91% of GABAergic neurons). In both protocols, a shift from neurogenesis to gliogenesis was observed after two months of organoid cultures (Fig. 1f-g). We did not observe dramatic differences in cell type and state distribution between different cell lines (Extended Data Fig. 1e, 2d).

**Figure 1.**
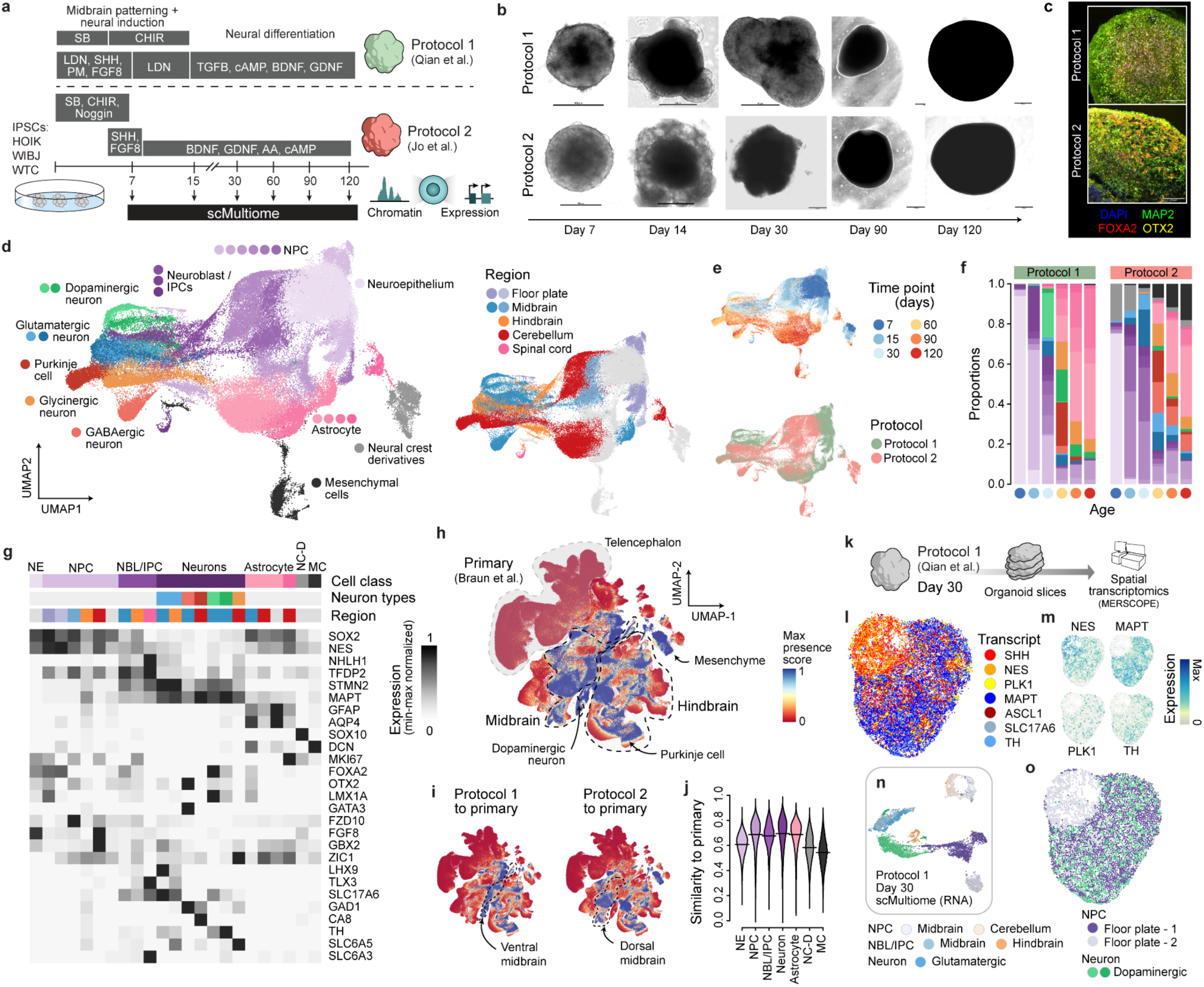
Single-cell atlas of human posterior neural organoid development. (a) Schematic of the two protocols generating midbrain/posterior brain organoids and the single-cell multi-omic measurements in time course. (b) Bright-field images of organoids showing examples of different stages of organoid development. (c) Immunofluorescence staining of MAP2, FOXA2 and OTX2 on organoids based on the two protocols. (d-e) UMAP of the posterior neural organoid developmental cell atlas based on gene expression profiles, colored by the annotation (d) of cell types (left) and regions (right), as well as experimental meta information (e) including time point (top) and protocol (bottom). NPC: neural progenitor cell. (f) Temporal cell type proportions profiles of organoids based on the two protocols. (g) Heat map of gene expression levels of selected cell type markers (row) across different cell types (columns). (h-i) UMAP of the human developing brain cell atlas colored by cell presence within the posterior neural organoid atlas (max presence scores), for both protocols together (h) or each of the two protocols separately (i). (j) Distributions of transcriptomic similarity between major cell classes in the posterior neural organoid atlas and their primary counterparts. (k) Schematic of spatial transcriptomic measurement on organoids based on Protocol 1 at day 30 with MERSCOPE. (l) Spatial distribution of selected transcript species on the example organoid section. (m) Expression levels of NES, MAPT, PLK1 and TH in the segmented cells on the example organoid section. (n) UMAP of a subset of the posterior neural organoid atlas, with the matched protocol and time point as the MERSCOPE data, colored by annotated cell types. NBL, neuroblast; IPC, intermediate progenitor cell. (o) Transferred cell type labels of the segmented cells on the example organoid section.

We next compared organoid data with a human first-trimester developing brain cell atlas^26^ using transfer learning^27^. Projection-based cell matching, as indicated by the maximum presence score per cell in the primary atlas (Fig. 1h-i) and transferred cell class and regional labels (Extended Data Fig. 2h-i), confirmed that cells produced with the two protocols matched to primary counterparts. While protocol 1 mapped to ventral midbrain fates, protocol 2 mapped to dorsal midbrain identities. Transcriptomic comparison between NPCs, neuroblast, intermediate progenitor cells (IPCs) and neurons in organoids showed high transcriptomic similarity to matched primary metacells (Fig. 1j). Interestingly, there were time-dependent patterns of transcriptomic similarity to primary neurons, with most neuron cell types peaking in similarity at 1-2 months of culture (Extended Data Fig. 2j). Differential expression (DE) analysis suggests that metabolism and synaptic programs may drive the divergence between neurons in organoids and those in the human developing brains (Extended Data Fig. 2k, Supplementary Table 2).

We performed spatial transcriptomics (MERSCOPE technology, Vizgen) to characterize tissue architecture of day 30 organoids (protocol 1) (Fig. 1k). We observed a patterned organization of two distinct regions demarcated by the expression of either *SHH* and *NES*, or *MAPT* and *TH* (Fig. 1l, m, Extended Data Fig. 3a-d, i). We annotated segmented cells through correlation-based label transfer using protocol and time point matched counterparts from the multi-omic cell atlas (Fig. 1m-n, Extended Data Fig. 3e-g). This revealed distinct spatial cell type compositions, where *SHH*+/*NES*+ regions were enriched for NPC floor plate type-2 cells, and *MAPT*+/*TH*+ regions were enriched for NPC floor plate type-1 and dopaminergic neurons. Notably, NPC floor plate type-1 cells are transcriptomically more similar to dopaminergic neurons, compared to the other floor plate NPCs, which implies the coordination of spatial patterning and neural differentiation.

### Chromatin accessibility dynamics during human posterior brain organoid development

We next analyzed chromatin accessibility dynamics in the multi-omic atlas of posterior brain organoid development. In total, 243,296 genomic regions were detected with significant coverage and region accessibility profiles show strong heterogeneity across the different time points and cell types (Fig. 2a-d; Extended Data Fig.4a-c). We performed two parallel analyses to identify gene regulatory regions underlying cell type differentiation. First, we used cisTopic to perform topic analysis on accessibility profiles^28^, identifying 49 clusters of co-accessible regions (regulatory topics) (Fig. 2e). Second, we used a generalized linear model based differential accessibility test, and identified 74,299 regions with significantly differential accessibility across cell types (DA regions). Notably, 21 of 49 regulatory topics have significant DA across cell types (one-sided Fisher’s exact test, bonferroni-corrected p-value<0.01) (Fig. 2g-h). Interestingly, we found distinct sets of DA regions marking different neuron cell types, but did not observe pan-neuronal accessible region clusters, implying that the genome-wide chromatin accessibility dynamics are critical in specification of distinct neuron cell type. This analysis, coupled with differential expression analysis for the downstream genes, also allows the identification of putative regulatory elements defining the identity of different neuronal cell types (Fig. 2i, Extended Data Fig. 4d-e, Supplementary Table 3, 4).

**Figure 2.**
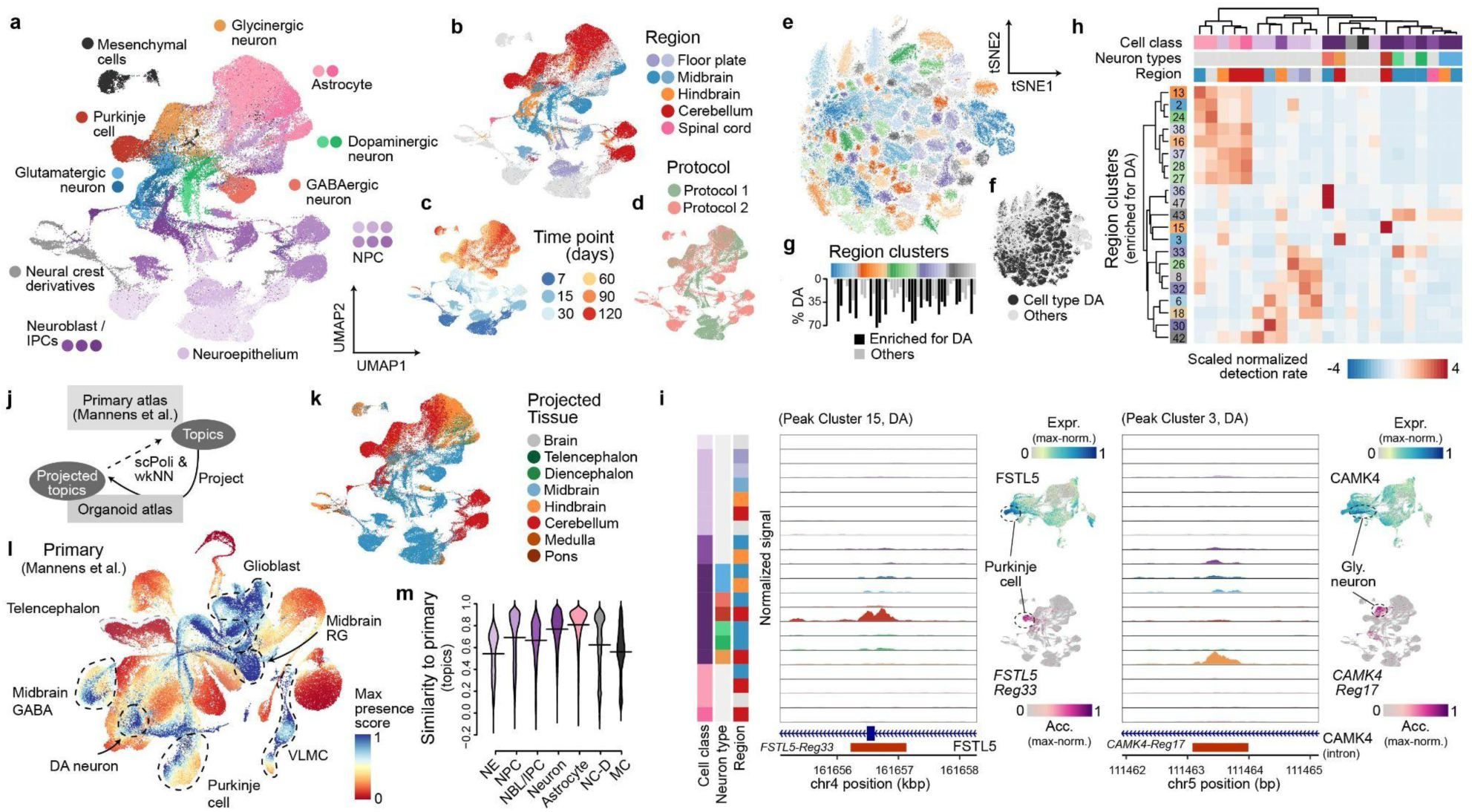
Chromatin accessibility dynamics during human posterior brain organoid development. (a-d) UMAP of the human posterior brain organoid cell atlas, based on chromatin accessibility profiles, colored by annotated cell type (a), regional identity (b), time point (c), and protocol (d). (e-f) UMAP of chromatin accessible regions, colored by the identified region clusters (e) and whether regions are differentially accessible (DA) in cell types (f). (g) Proportions of DA regions in region clusters. (h) Heat map shows average relative accessibility patterns across cell types (columns) for different region clusters (rows). (i) Chromatin accessibility profile tracks in different cell types at two representative loci: FSTL5 and CAMK4. (j) Schematic of topic-based projection to the primary human developing brain scATAC-seq cell atlas. (k) UMAP of the human posterior brain organoid cell atlas, colored by transferred tissue labels from the primary cell atlas. (l) UMAP of the human developing brain cell atlas colored by cell presence within the posterior neural organoid atlas (max presence scores). (m) Distribution of chromatin accessibility topic profile similarity between major cell classes in the posterior neural organoid atlas and their primary counterparts.

To benchmark how well organoid chromatin accessibility dynamics recapitulate human early brain development, we developed an approach combining topic analysis and transfer learning^29,27^ to project organoid scATAC-seq data to a single-cell chromatin accessibility atlas of the human developing brain^30^ (Fig. 2j). This allows estimation of which cells of the primary reference are prevalent in organoids, as well as estimation of similarity in chromatin accessibility between organoid and matched reference cells. Organoid cells were matched to primary counterparts, with NPCs, neuroblast, IPCs and neurons showing high similarities in chromatin accessibility topics (Fig. 2k-m, Extended Data Fig. 4f–i). Transferred cell class and region labels based on transcriptome or chromatin accessibility profiles are largely consistent, indicating that cells in the organoids recapitulate major aspects of transcriptome and chromatin features of primary brain development.

### Inferring and perturbing regulatory programs of human posterior brain organoid development

We used gene regulatory network (GRN) inference to understand how differentiation and specification of different cell types are regulated in human posterior brain organoids^31^. We globally modeled the interaction between transcription factor (TF) expression, binding motif accessibility, and target gene expression. We identified a GRN involving 344 TFs, accounting for 342 positive (activating) TF regulons and 258 negative (repressive) TF regulons, largely reflecting cell class differences (Fig. 3a-b; Extended Data Fig. 5a-d). Regulons showed significant activity differences between cells with midbrain and hindbrain identities, while NPCs and neurons within a given region showed correlated regulon activities, TF expression and motif enrichment (Fig. 3c-d, Extended Data Fig. 5e). We identified TFs with regulon activities that correlate with neuron cell type identities including *LHX1* for Purkinje cells, *OTX2* for GABAergic midbrain neurons, *FOXA2* for dopaminergic neurons (Fig. 3e-g, Extended Data Fig. 5f).

**Figure 3.**
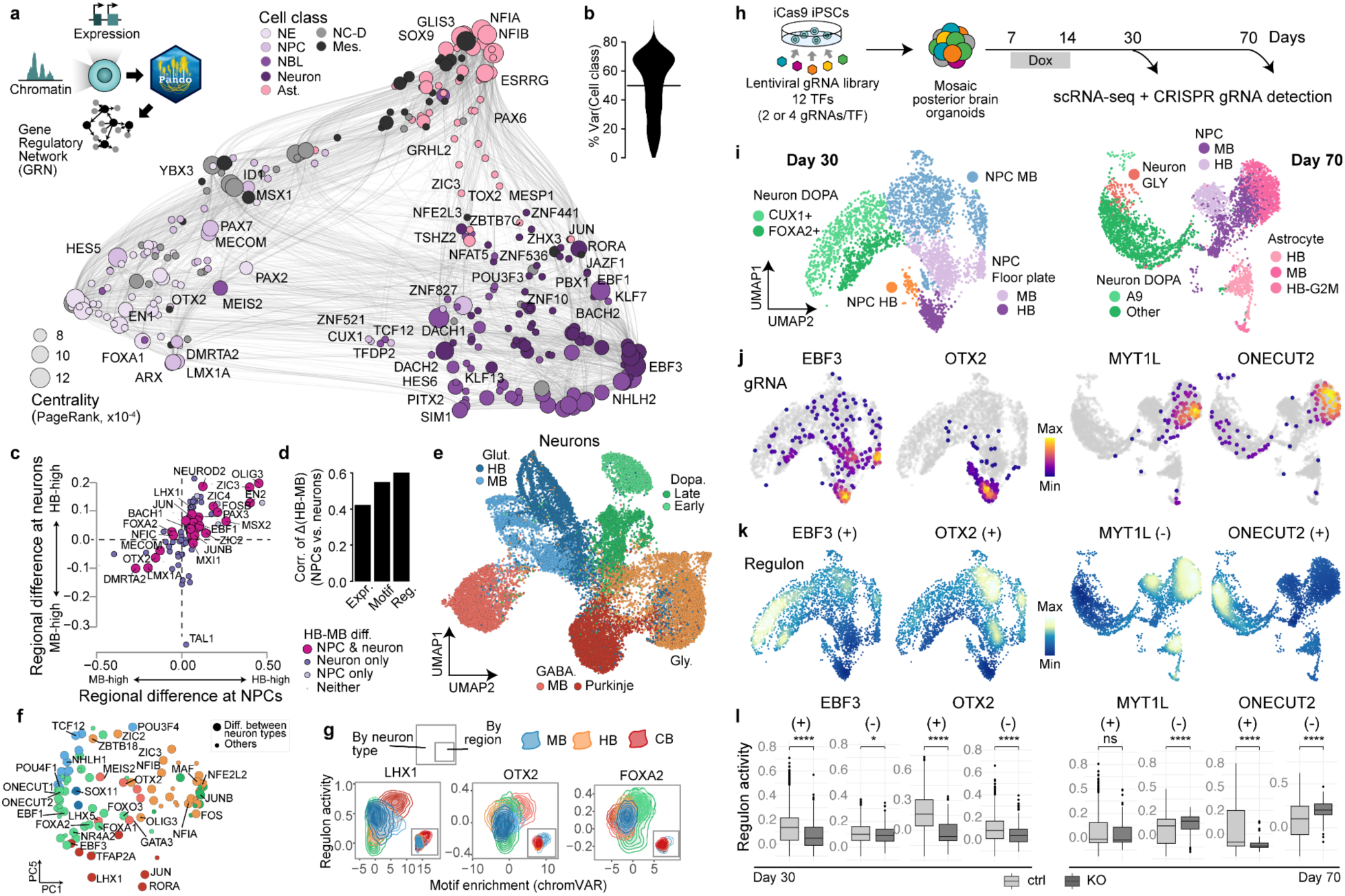
Inferring and perturbing human posterior brain organoid developmental regulomes. (a) Schematic of gene regulatory network (GRN) inference using Pando framework (top left); UMAP embedding of the inferred TF network based on co-expression and inferred interaction strength between TFs. Color of dots represents the cell class with the highest expression value of the indicated TF, size of dots represents PageRank centrality of each TF. (b) Violin plot showing the variation of module activity, explained by cell class. (c) Comparison of regional differences of the regulon activity for neurons (y-axis) and for NPC (x-axis). Regulon activities of NPC and neurons were summed up based on regional identity and then the difference between hindbrain and midbrain was calculated for each TF. The size and color of the dots represent the results of the analysis of the differential regulon activity between midbrain and hindbrain: if the regulon was differentially active for both neuron and NPC, only in one of them, or in neither. (d) Barplot showing correlation of hindbrain-midbrain differences of TF expression, motif enrichment and regulon activities between NPC and neurons. (e) UMAP embeddings of 24,153 neuronal cells, constructed based on weighted multimodal neighbours graph, representing a weighted combination of RNA and ATAC-seq modalities. Colors represent neuronal subtypes. (f) PCA embedding of selected TFs, coloured by neuron subtype with the highest expression value of each TF. The dot size indicates whether regulon is classified as differentially active between neuron subtypes. (g) Density plot, indicating motif enrichment and regulon activity for selected transcription factors for each neuron subtype. The small panel in the right colour represents the same analysis, split by region identity of neurons. (h) Schematic, illustrated single-cell pooled TF knock-out (KO) experiment. (i-k) UMAP embeddings of the cells with detected gRNA for day 30 (left) and day 70 (right) time point, colored by cell type identity (i), detected gRNA density (j), and positive (+) or negative (-) regulon activity for each indicated TF. (k). (l) Barplots, showing the difference in positive (+) and negative (-) regulon activity between control and KO cells for day 30 organoids (left) and day 70 organoids (right). *** p-value < 0.001, ** 0.001< p-value < 0.01, * 0.01 < p-value < 0.05.

We used a pooled, multiplexed CRISPR/Cas9 gene knock-out (KO) experiment with single-cell transcriptomic readout to perturb and validate selected TF regulomes (Fig. 3h). We designed gRNAs and generated a pooled lentiviral library targeting 12 TFs with specific midbrain or hindbrain expression or high regulatory centrality (Extended Data Fig. 6a-b; Supplementary Table 5). IPSCs carrying a doxycycline-inducible *Cas9* cassette in the AAVS1 safe harbor locus were transduced, mosaic organoids were generated, and perturbations were induced at neuroepithelium stage from day 7 to 14. Mosaic organoids were analyzed using scRNA-seq and gRNA amplicon sequencing at day 30 and day 70, recovering 31,857 cells, among which a gRNA was detected in 8711 cells (Fig. 3i, Extended Data Fig. 6c-d).

We observed differential cell type abundance associated with specific perturbations in day 30 organoids. For example, perturbation of *OTX2*, a well-known midbrain fate regulator^32^, resulted in complete shift in cell type composition towards hindbrain regions (Fig.3j, Extended Data Fig. 6e). *OTX2* regulon activity was highest in midbrain cells (Fig.3k), with the regulon suppression activity in KO cells (Fig.3l). As another example, *EBF3* perturbation resulted in an enrichment of NPCs, consistent with regulon activity in neurons (Fig. 3j-k). *EBF3* is inferred to regulate the expression of numerous neural differentiation genes, including *MYT1L, ONECUT1,* and *ONECUT2* (Extended Data Fig.6e-h, Supplementary Table 6). DE gene analysis between *EBF3*-KO and control cells revealed that downregulated genes were enriched in synapse organization, assembly, and membrane repolarization functions (Extended Data Fig.6h). This finding aligns with the shift in cell type composition observations, and suggests that *EBF3* plays an important role in the transition from NPC to neuron by activating key neural differentiation genes, possibly explaining developmental disruption resulting from EBF3 mutation^33^.

At day 70 of organoid development, we observed the increased prevalence of *MYT1L*-KO cells in astrocytes, which is in concordance with the well known role of this gene in neurogenesis, suppressing the differentiation of non-neuronal fates (Fig. 3j)^34^. Furthermore, we have observed increased activity of *MYT1L* negative regulons in the KO cells (Fig. 3k-l), which aligns with *MYT1L* acting primarily as a repressor^35^. *ONECUT2* is another gene where perturbation disrupts NPC and neuron abundance (Fig. 3j-k), with perturbed cells depleted in neurons and enriched in astrocytes. Genes upregulated in *ONECUT2*-KO cells are related to gliogenesis, whereas downregulated genes are related to synapse transmission, neurotransmitter transport and secretion (Extended Data Fig.6i-l, Supplementary Table 6), which is consistent with the role of *ONECUT2* as a neural fate inductor^36^. Together, these data provide a multiplexed single-cell perturbation experiment in organoids to examine the effects of genetic perturbation on posterior cell fate specification, suggesting mechanisms underlying gene functions.

### Combinatorial morphogen screen introduces a spectrum of organoid models of the posterior human brain

Posterior cell type diversity is underrepresented in current brain organoid protocols^21^. We devised a screen to systematically evaluate if morphogen combinations can induce a new spectrum of posterior neuron subtypes (Fig. 4a). We based the screen on protocol 1 including neural induction using dual SMAD inhibition during the first 7 days of organoid culture, and varied exposure to morphogen combinations in different time windows with a major focus on the second week of organoid development during initial neuronal differentiation (Fig. 4a). We chose known modulators of anterior-posterior (AP) patterning (WNT signaling agonist CHIR-99021, CHIR; Retinoic acid, RA; Fibroblast growth factors (FGF) FGF-8, FGF-2, and FGF-17; R-spondin 2 and 3) as well as dorso-ventral patterning (Bone morphogenetic protein BMP4; Insulin; Sonic hedgehog, SHH; SHH pathway agonist purmorphamine, PM) and designed 48 distinctive morphogen modulator conditions across a range of concentrations and combinations (Supplementary Table 7).

**Figure 4.**
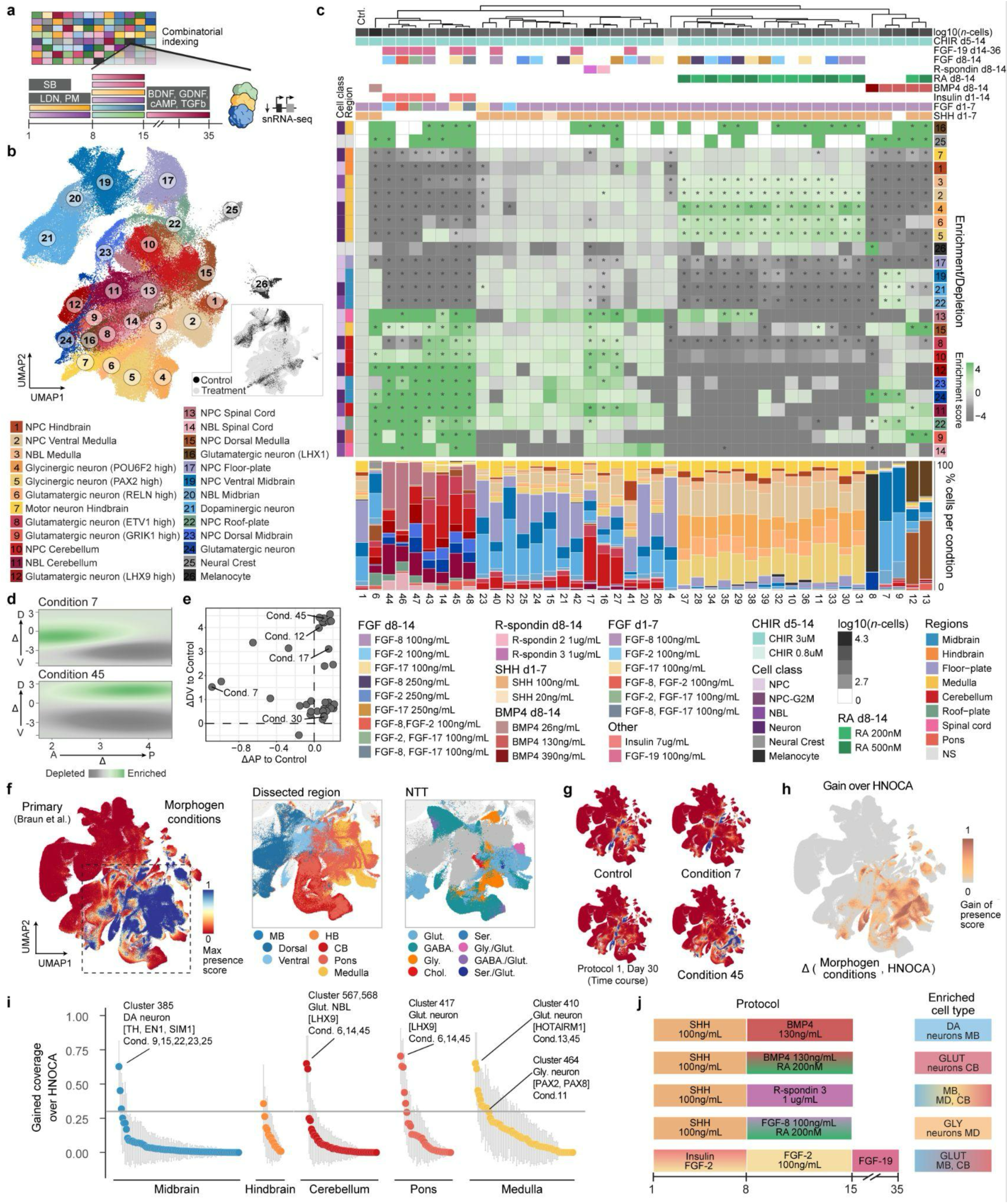
A combinatorial morphogen screen expands posterior organoid protocols. (a) Experimental scheme and timeline of the morphogen combinatorial patterning screen with single-cell transcriptomics readout. 96 individual organoids exposed to each of the 43 morphogen combinations were individually analyzed with snRNA-seq using split-pool combinatorial barcoding (Parse Biosciences). (b) UMAP embedding of 177,718 cells in the dataset, colored by cluster identity (left) and mode of treatment (bottom right). NBL, neuroblast; NPC, neural progenitor cell. (c) Overview of conditions and results of combinatorial morphogen screen. Heatmap showing enrichment/depletion scores (log2 of cell types proportions in relation to control), scores with p-value <0.05 (FDR adjusted using Benjamini-Hochberg procedure) marked with *. Each column represents a distinct morphogen combination with color-coded annotation on the top of the column. Each row corresponds to a cluster with color-coded annotation: along the left side - Cell class and Region, along the right side - cell types, corresponding to the ones, presented in the UMAP embeddings in panel (b). Barplots (bottom) showing the relative composition of organoids per each condition, condition names are labeled in the bottom. Morphogen conditions and cell types were hierarchical clustered based on enrichment/depletion values, the control condition (label 1) was not included in the clustering procedure and was added later for visualization purposes. NS, non-specific; Ctrl.,control. (d) Visualisation of difference of spatial mapping of selected morphogen treatment conditions in comparison to control using the dorso-ventral (vertical) and rostro-caudal (horizontal) axis of the developing human brain radial glia^26^. D, dorsal; V, ventral; A, anterior; P, posterior. (e) Scatterplot showing aggregated location of morphogen screen condition along DV and AP axis in relation to control. Cond., condition. (f) UMAP of the primary reference, colored by the max presence scores across all screen conditions. The area, indicated with black line, is further colored by dissected regions (center) and neural transmitter transporter (right). MB, midbrain; HB, hindbrain; CB, cerebellum; NTT, neural transmitter transporter; Glut., glutamatergic; GABA., gabaergic; GLY., glycinergic; Chol., cholinergic; Ser., serotonergic. (g) UMAP of the primary reference, colored by the max presence scores across the indicated screen conditions and protocol 1 day 30 time point from posterior brain organoid atlas. (h) UMAP of the primary reference, colored by gained coverage of posterior brain organoid morphogen screen over human neural organoid cell atlas (HNOCA)^21^ with negative values trimmed to zero. (i) Gain of cell cluster coverage of the screen conditions over to HNOCA, corresponding to average max presence scores per primary reference cluster with negative values trimmed to zero. The threshold is set to 0.3 and represented by a gray horizontal line. Circles show mean cluster gain and coloured by the most abundant dissected region of the cluster, gray vertical lines indicate standard deviation of gain coverage. DA, dopaminergic. (j) Schematic, showing the protocols, tested in the morphogen screen, allowing for enrichment of the representative cell types.

Organoids were grown under 48 different morphogen conditions, differentiated until week 5 and individually dissociated for single-nucleus RNA-seq profiling using combinatorial barcoding^37^. Notably, 5 of the 48 conditions resulted in organoids with reduced size and were excluded from snRNA-seq profiling. In total, 177,718 high-quality cells were profiled across 43 conditions, with a median of 2,798 cells and 2 individual organoid replicates per condition (Extended Data Fig. 7a-b). Clustering of the single-cell transcriptomic data revealed 25 molecularly distinct cell types, with strong cell type abundance changes dependent on morphogen treatments, particularly for RA and BMP4 (Fig. 4b, Extended Data Fig. 7c). Cell type composition across individual organoids for a given condition was highly reproducible (Extended Data Fig. 7d)

Based on organoid cell type composition, we classified morphogen effects into at least five groups (Fig. 4c, Supplementary Table 8): Lack of SHH or Insulin generated glutamatergic neurons from midbrain, cerebellum and pons; BMP4 inclusion at an intermediate dose resulted in an increased proportion of ventral midbrain progenitors and dopaminergic neurons (clusters 20, 21)^38,39^; RA conditions resulted in a variety of ventral medulla cell types (clusters 1-7), including two subtypes of glycinergic neurons; the combination of BMP4 and RA led to the emergence of neuronal cells of the dorsal medulla (clusters 15,16); and both R-spondins, as well as BMP4 at a low dose, gave rise to a diverse range of midbrain, cerebellum and medulla cell types that were shown using shannon diversity index calculation (Extended Data Fig. 7e) to represent the conditions with highest cell type diversity.

Significant shifts in the composition of dopaminergic (DA) neuron subtypes were also observed in treatment conditions in comparison to the control (Extended Data Fig. 7f-h). The administration of R-spondin 3 and a low dose of BMP4 resulted in an increased proportion of *FBN2*+ DA subtypes, whereas FGF-2 in low dose (100 ng/mL), provided early and late, resulted in the increased proportion of A9-like DA neurons. The administration of BMP4 at an intermediate dose alone increased the proportion of *SOX6*+, *OTX2*+ DA neurons, while its combination with RA led to the emergence of a greater proportion of *SLC17A6*+ (*Vglut2+*) DA neurons in a RA dose-dependent manner.

To quantify the influence of screened morphogen conditions on dorso-ventral (DV) and antero-posterior (AP) fate identity, we calculated DV and AP scores for each sequenced organoid progenitor cell using a regularized linear model, trained on the radial glia of the human developing brain cell atlas^26^ (methods). Subsequently, for all progenitors in a given treatment condition, the differential AP and DV scores relative to the control were calculated (Fig. 4d-e, Extended Data Fig. 8a-c). Condition 7, for instance, exhibits a slight dorsal and anterior shift relative to the control, resulting in a high proportion of midbrain dopaminergic neurons. In contrast, condition 45 displays a dorsal and posterior shift, predominantly comprising cell types with cerebellar identity.

To assess correspondence of the organoid cells generated under the different morphogen conditions to human primary brain cell types or states, we mapped the morphogen screening data to a primary human developing brain cell atlas^26^ using transfer learning^27^, as described above. We estimated presence scores for every primary cell type in each morphogen screen condition (Fig. 7f) ^21^ and found that screen conditions were enriched for midbrain and hindbrain region identities, while telencephalic or diencephalic cell types were absent (Fig. 7f, Extended Data Fig.8d-f). Notably, the presence scores of control organoids from the screen are consistent with those of organoids from our previous time course dataset (Fig. 7g), indicating robustness of the organoid protocol.

We next evaluated whether the morphogen screen conditions generated cell states previously absent or underrepresented in *in vitro* organoid protocols. We therefore compared the max presence scores of the morphogen screen data conditions to those estimated for the integrated human neural cell organoid atlas (HNOCA)^21^. This analysis revealed that cell types from midbrain and hindbrain regions were enriched in the morphogen screen (Fig. 4h), with a total of 28 reference cell clusters showing a significant abundance increase in certain treatment conditions (Fig. 4i, Extended Data Fig.8g). The most enriched clusters included dopaminergic neurons from the ventral midbrain, *LHX9*+ glutamatergic neurons in cerebellum and pons, and *PAX2*+/*PAX8*+ glycinergic neurons in medulla.

In summary, our morphogen screen allows the expansion of the diversity of posterior brain cell types generated with *in vitro* organoid cultures and serves as a rich resource for the community to fine-tune organoid protocols (Fig. 4j).

### Differential abundance-informed trajectory inference provides inroads to understand human brain development

The shared differential abundance of progenitor and neuronal populations across morphogen patterning conditions can inform their relation within a differentiation trajectory (Fig. 5a). We therefore sought to identify progenitor-neuron differentiation trajectories in the morphogen screening data informed by the detected differential abundances. We calculated pairwise Pearson correlation (*r*) between proportions of cell types across conditions, and hierarchically clustered cell types using the correlation distance (defined as 1-*r*) matrix (Fig. 5b). This allowed the identification of nine cell type clusters, which could be summarized into five distinct NPC-neuron trajectories (Fig. 5c). Importantly, the annotated regional identity of NPC and neuron populations assigned to the same trajectory matched, such as ventral midbrain NPCs, neuroblasts and dopaminergic neurons, assigned to the ventral midbrain trajectory, or ventral medulla NPCs, medulla neuroblasts, glycinergic neurons, and *RELN*-high glutamatergic neurons assigned to the medulla trajectory. This further supports the validity of the inferred trajectories. Notably, while the shared differential abundance of NPC populations can be positively correlated with their transcriptomic similarities (Extended Data Fig. 9b), there are cases where NPC populations with high transcriptomic similarity are exclusively found across conditions, indicating that transcriptomic similarity can mislead trajectory inference.

**Figure 5.**
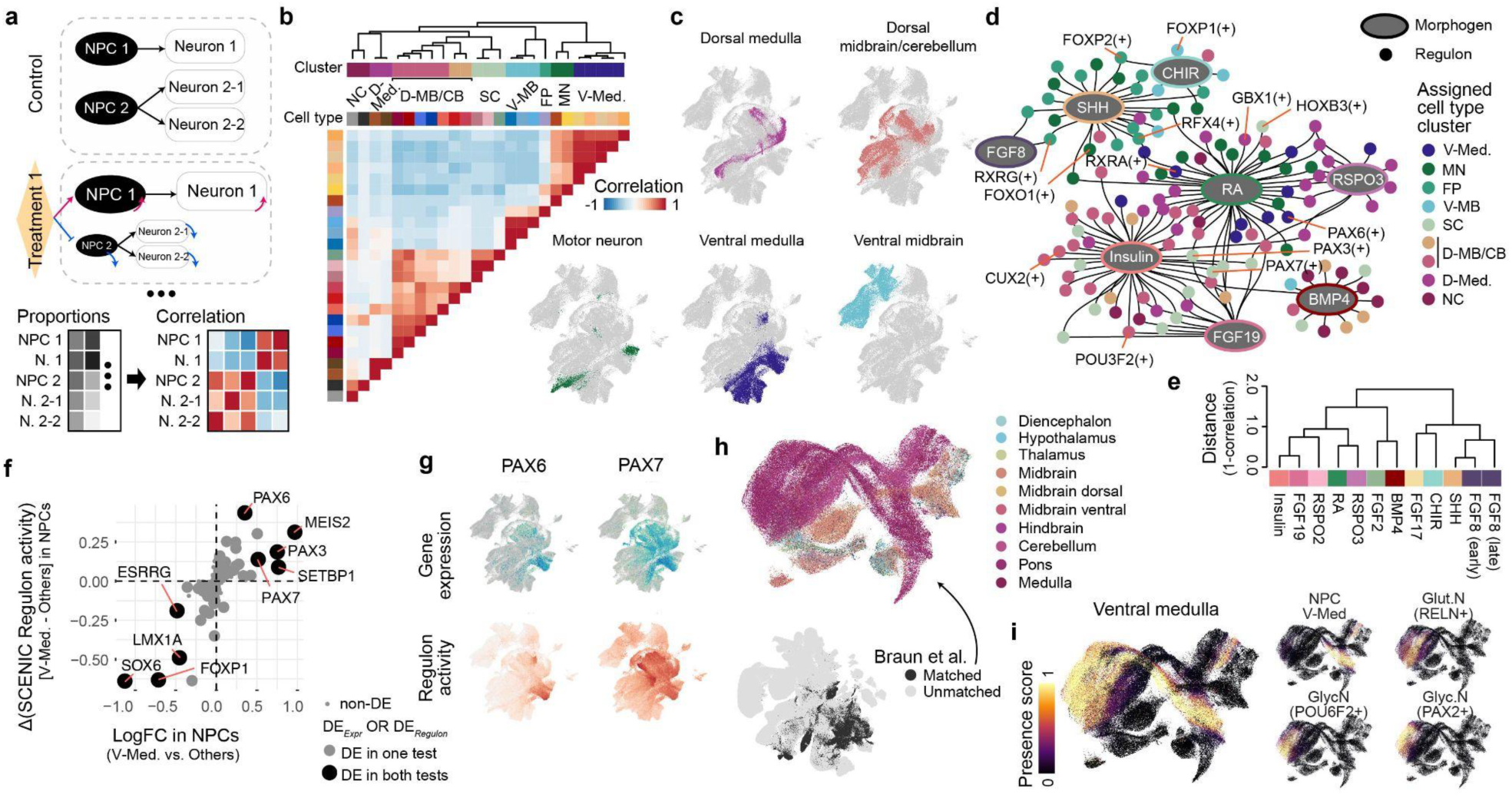
Perturbation-informed trajectory inference reveals regulatory mechanisms underlying neural differentiation trajectories. (a) Schematic of differentiation trajectory inference based on cell type compositions in multi-treatment-control experiments. (b) Hierarchical clustering of cell types in the morphogen screening experiments, based on the pairwise correlation (shown by the heat map) of cell type proportions in different morphogen conditions. (c) UMAPs show five differentiation trajectories of different neuron cell types revealed by the composition-based cell type clusters. (d) Morphogen-regulon regulatory relationship with GRNBoost2. (e) Hierarchical clustering of morphogen treatments, based on their importances in activating/inhibiting regulon activities. (f) Differential expression of transcription factors (TFs) and differential activity of their SCENIC-reconstructed regulons, between ventral medulla neural progenitor cells (NPCs) and other NPCs in the morphogen screening data. (g) UMAPs show expression and regulon activities of two example TFs, PAX6 and PAX7, in the morphogen screening data. (h) UMAP of the human developing brain cell atlas. The bottom one shows whether a cell is matched to any cell type in the morphogen screening data (the matched primary sub-atlas). The upper panel shows the UMAP of the matched primary sub-atlas, colored by the dissected brain regions.. (i) UMAP of the matched primary sub-atlas, colored by presence within the ventral medulla cell clusters in the morphogen screening data.

To explore the gene regulatory logic underlying the diversification of NPC pools in the presence of morphogen cues, we performed a hierarchical regulatory network reconstruction analysis as previously described^40^. Briefly, we identified TF regulons in the screening data using SCENIC^41^, (Extended Data Fig. 9c-e) and inferred their regulatory relationship with morphogens using GRNBoost2^42^. This analysis linked morphogens to different sets of regulons that showed distinct activity patterns along the different neuronal differentiation trajectories (Fig. 5d). For instance, RA was linked to many regulons with enriched activities in the ventral medulla (e.g. *PAX6*(+), *RXRA*(+)) and dorsal medulla (e.g. *GBX1*(+)) trajectories, indicating the essential role of RA in activating medulla-specific genetic programs. Some regulons showed linkage to multiple morphogens, such as *PAX7*(+), which was activated by both RA and insulin. Based on the inferred regulatory weights and directions for morphogens on different regulons, we were also able to evaluate the similarities of different morphogens’ influence on neuron cell fate determinations (Fig. 5e). Focusing on the ventral medulla trajectory, we identified *PAX6* and *PAX7* as critical fate regulators linked to RA and with enriched expression and regulon activities in ventral medulla NPCs compared to other NPC populations (Fig. 5f, g). In addition, TFs such as *SOX6*, *FOXP1* and *LMX1A* showed negative correlation with ventral medulla identity, suggesting that these TFs and their regulons need to be turned off in order to adopt ventral medulla fate.

To assess the *in vivo* relevance of the identified differentiation trajectories, we subsetted the human developing brain cell atlas^26^ to cells confidently mapping to the organoid morphogen screening data (Fig. 5h, Extended Data Fig. 8d). Focusing on primary NPCs and neurons matching the organoid ventral medulla cells (Extended Data Fig. 9i), we could reconstruct a trajectory from NPCs towards three different neuronal populations: *RELN*-high glutamatergic neurons, *POU6F2*-high glycinergic neurons and *PAX2*-high glycinergic neurons (Fig. 5i, Extended Data Fig. 9h), consistent with the organoid ventral medulla trajectory. Pseudotemporal gene expression dynamics in the organoid data showed high similarity to the primary ventral medulla trajectory (Extended Data Fig. 9j-k). Finally, we compared the performance of differential abundance-informed trajectory inference to that of conventional transcriptome-similarity based trajectory inference methods ^43^. While certain regional differentiation trajectories could be readily inferred from the primary brain atlas data using transcriptome-similarity alone (e.g. dorsal midbrain), others failed, likely due to high transcriptomic similarity to progenitor populations from other brain regions (e.g. dorsal medulla) (Extended Data Fig. 9l-m). In those cases, information about differential abundance in the organoid screening data provided important additional information to resolve trajectories. Altogether, these data highlight that systematic *in vitro* organoid morphogen perturbations enable robust trajectory inference across brain regions.

## Discussion

Here, we present a thorough single-cell multi-omic atlas of human posterior brain organoid development. We utilized two existing protocols originally developed to generate midbrain cell types and found that both protocols, in addition to midbrain identities, generate significant portions of hindbrain cells. Similar observations were made for other midbrain protocols^44,45^, indicating that further efforts are required to achieve higher regional specificity in *in vitro* cultures. Furthermore, it highlights the importance of organoid protocol profiling at the single-cell level for comprehensive benchmarking and validation of cell type diversity.

To provide insights into the mechanisms underlying human brain regionalization, we complemented the single-cell multi-omic atlas with genetic perturbation datasets. We have identified brain region-specific programs, many of which include well-known master-regulators, such as *OTX2, LMX1A, LHX1, EN2, ZIC3*. Interestingly, the majority of programs are shared between NPCs and neurons on the level of regulon activity, TF expression and motif enrichment, implying that similar genetic programs are responsible for regional identity establishment and maintenance. It provides evidence for a deterministic specification of neural diversity^46^.

We use systematic modulation of morphogen signalling pathways to expand the diversity of *in vitro*-generated cell types of the posterior human brain. The approach generated underrepresented brain cell types and identified protocols that yield cerebellum- and pons-specific *LHX9*+ glutamatergic neurons as well as glutamatergic and two types of glycinergic neurons from the dorsal and ventral medulla, respectively. The screen shows how combinations of morphogen modulators influence the regional identity along the dorsal-ventral and anterior-posterior axes, as illustrated by BMP4 and RA. BMP4 alone has the dorsalising effect on the tissue, resulting in the enrichment of *Vglut2+* DA neurons, RA alone stimulates the production of posterior ventral cell types, while the combination of RA and BMP4 results in generating dorsal posterior cells. Notably, the observed effects of RA were dominant, overriding the effects of various FGFs. This suggests to test milder caudalization agents, such as CHIR or lower concentrations of RA, to achieve a higher degree of granularity in brain stem patterning.

We incorporate differential cell type abundance into trajectory inference and reconstruct differentiation events within the morphogen screen data. As an example, dorsal and ventral medulla trajectories were exclusive in different conditions in the morphogen screen data, however dorsal and ventral medulla NPCs showed high transcriptomic similarity in both the screen data and the primary atlas, making it difficult to resolve differentiation trajectories purely using transcriptomic information. This highlights how organoid morphogen screening data can inform the reconstruction of differentiation trajectories in primary atlases in cases where transcriptome-based methods fail.

Altogether our data demonstrates the potential of organoids to recapitulate primary human posterior brain development at the transcriptomic and chromatin level. Furthermore, it reveals an extraordinary diversity of cell types in the posterior brain, providing an integrated overview of morphogen-induced regionalization of posterior brain organoids. The data demonstrates that the morphogen screens with single-cell genomic readouts present novel opportunities to optimize new region-specific protocols as well as open up new ways of exploring neuronal differentiation trajectories.

## Methods

### Experimental methods

#### Stem cell and organoid culture

We used three human iPS cell lines (HOIK1, WIBJ2 from the HipSci resource^43^ and WTC from the Coriell Institute). Stem cell lines were cultured in mTeSR Plus with mTeSR Plus supplement ((Stem Cell Technologies, 100-0276) and supplemented with penicillin–streptomycin (1:200, Gibco, 15140122) on Matrigel-coated plates (Corning, 354277). After splitting with TryplE (Gibco, 12605010) or EDTA in DPBS (final concentration 0.5 mM) (Gibco, 12605010) cells were passaged 1–2 times per week. Rho-associated protein kinase (ROCK) inhibitor Y-27632 (final concentration 5 μM, STEMCELL Technologies, 72302) was provided the first day after passage. Cells were tested for mycoplasma infection regularly using PCR validation (Venor GeM Classic, Minerva Biolabs) and found to be negative. A total of 10,000 cells were plated in ultra low-attachment plates (Corning, CLS7007) to generate brain organoids using midbrain organoid differentiation protocols^13,14^.

#### Single-cell multiome experiments for the developmental time course

Brain organoids were generated from three different stem cell lines (WTC, WIBJ2, HOIK1) simultaneously. Brain organoids of the same batch were dissociated at multiple time points of the course of brain organoids development: neural induction (day 7) and neural differentiation and maturation (days 15, 30, 60, 90, 120). Organoids of the three different cell lines were pooled on the basis of size and dissociated together, and the cell lines were later demultiplexed on the basis of the single-nucleotide polymorphism information. Multiple organoids of each line were pooled together to obtain a sufficient number of cells. If needed, on the later time points organoids were cut in halves and washed three times with HBSS without Ca2+ and Mg2+ (STEMCELL Technologies, 37250). Tissue dissociation to single cell suspension was done with a papain-based dissociation kit (Miltenyi Biotec, 130-092-628). Prewarmed papain solution (2 ml) was added to the organoids and incubated for 15 min at 37 °C. This was followed by enzyme mix A addition and then tissue pieces were triturated 5–10 times with 1,000 μl wide-bore and P1000 pipette tips. After that the tissue pieces were incubated twice for 10 min at 37 °C with trituration steps with P1000 and P200 pipette tips. After dissociation cells were filtered with 30 um filters and centrifuged. Before nuclei isolation, 100,000 cells were washed twice with 50 μL PBS containing 0.04% BSA. To isolate nuclei from the ready single-cell suspension, cells were resuspended in 50 μL lysis buffer (10 mM Tris-Cl pH7.4, 10 mM NaCl, 3 mM MgCl_2_, 1% BSA, 0.1% Tween-20, 1 mM DTT, 1U/μL RNase inhibitor (Roche Protector RNase-Inhibitor), 0.1% NP-40, 0.01% Digitonin (Invitrogen, BN2006)) and incubated for 3 minutes on ice, neutralized by adding 50 uL wash buffer (10 mM Tris-Cl pH7.4, 3 mM NaCl, 10 mM MgCl2, 1% BSA, 0.1% Tween-20, 1 mM DTT, 1U/μL RNase inhibitor). Single-nuclei suspension-generation and library preparation were performed according to the 10x Chromium Single Cell Multiome ATAC + Gene Expression kit protocol.

#### Immunohistochemistry

Organoids were fixed in 4% PFA at 4 °C overnight, followed by incubation in 30% sucrose solution for 24–48 h. Afterwards, organoids were transferred to plastic cryomolds (Tissue Tek) and embedded in OCT compound 4583 (Tissue Tek) for snap-freezing on dry ice. For immunohistochemical stainings, organoids were sectioned in slices of 20 μm thickness using a cryostat (Thermo Fisher Scientific, Cryostar NX50). Organoid sections were quickly washed in PBS to remove any residual OCT. The sections were then incubated in antigen-retrieval solution (HistoVT One, Nacalai Tesque) at 70 °C for 20 min. Excess solution was washed away with PBS and the tissue was incubated in blocking-permeabilizing solution (0.3% Triton X-100, 0.2% Tween-20 and 5% normal donkey serum in PBS) for 1 h at room temperature. Next, the sections were incubated overnight at 4 °C in blocking-permeabilizing solution containing rabbit anti-MAP2 (1:1000, Sigma-Aldrich, AB5622), mouse anti-OTX2 (1:200, Invitrogen, MA5-15854) and goat anti-FOXA2 (1:200, R&D systems, AF2400) antibodies. The next day, the sections were rinsed three times in PBS before incubation for 1 h at room temperature with 1:500 secondary antibody (donkey anti-rabbit Alexa 488, ab150073 and donkey anti-mouse Alexa 568, ab175472 and donkey anti-goat Alexa 647, ab150135) in blocking-permeabilizing solution. Finally, the secondary antibody solution was washed off with PBS and the sections were stained with DAPI before covering with ProLong Gold Antifade Mountant medium (Thermo Fisher Scientific). Stained organoid cryosections were imaged using a confocal laser scanning microscope, and six different z-plane images (z-step = 2–3 μm) were acquired using a ×20 magnification objective. The images were further processed using Fiji.

#### Spatial transcriptomics experiment

To design the 500-gene panel for the spatial transcriptomic measurement with MERSCOPE, we combined a pre-defined gene list which includes canonical regional and cell type markers, and an additional list of genes selected by geneBasis^47^ (Supplementary Table 9).

We curated two data sets for the gene panel selection. The first data used for the process include public scRNA-seq data sets of brain organoids with no more than one-month of culture^23,31,48–50^, together with the scRNA-seq portion of the time-course scMultiome data of day 7, 15 and 30 presented in this study (the early data set). All the included scRNA-seq data were merged, normalized, and scaled (for the 3000 highly variable genes). Principal component analysis (PCA) was applied to the scaled data for the first 20 principal components (PCs). CSS^51^ was applied for data integration on samples. Louvain clustering (resolution=2) was performed to identify cell clusters for early brain organoid cultures. In addition, a merged data set of two public scRNA-seq data sets^48,49^ of brain organoids older than one month (the late data set) was curated and processed (union of highly variable genes per data set and the merged data set; PCA on the scaled data for the top-20 PCs; CSS integration on samples; louvain clustering with resolution=3 on the CSS representation). Differentially expressed genes (DEGs) were identified for each cluster with the wilcoxauc() function in the presto package (adjusted P<0.01, FC>1.2, AUC>0.65, detection rate difference>20% and detection rate ratio>2; max 20 genes per cluster based on the AUC value). The DEGs were merged and examined in cluster pairs along the branch merging on the dendrogram of clusters based on their expression distances. Clusters with no more than five overall DEGs showing DE were merged, resulting in the final clustering results for the late data set. Average gene expression profiles were estimated for each cluster in the early and late data sets in the unit of transcripts per million (TPM). For the pre-defined gene list, genes are required to satisfy one of the following criterias: 1) with the maximal average expression in the early data set clusters between TPM 1 and 500; 2) with maximal average expression between TPM 500 and 1000 but are not clustered with any genes selected by criteria 1; 3) with maximal average expression in the late data set clusters between TPM 1 and 500. This resulted in 242 genes in total.

Next, we used geneBasis to complete the 500-gene panel, using the early data set. To reduce the computational complexity, we reconstructed metacells for each sample separately using the approach described previously^21,48^, given the global PCs for neighbor graph construction and a downscaling ratio of 1:10, to pool the gene counts of cells from the same samples which are similar on the transcriptomic level together followed by normalization to the total transcript counts per metacell. The normalized expression of metacells were then provided to geneBasis to target for 600 genes in total, given the pre-selected genes mentioned above and a gene list to drop from the selection (the union of genes with maximal average expression in early data set cluster < 1 TPM and those with > 500 TPM, cell cycle related genes, mitochondrial genes and ribosomal protein genes). The resulting gene list was further filtered by the Vizgen database, and the top-500 genes remained were chosen for the spatial transcriptomics experiment by MERSCOPE.

Two organoids based on protocol 1^25^ at day-30 of their culture were fixed in 4% PFA for 1.5 hours and subsequently exposed to sucrose gradients (15% and 30%) until the organoids sank. Organoids were then co-embedded and frozen in an OCT (Tissue-Tek® O.C.T. Compound, Sakura) block before being stored at -80C. To increase the adherence of organoid slices to the MERSCOPE slides (Vizgen, #10500001), slides were coated with 1mL of Poly D-Lysine (Thermo Scientific #A3890401) for 3 hours and washed with 2mL RNAase free water three times. Slides were dried and left at room temperature. The block was incubated for 30 minutes at -20°C in a cryostat (ThermoFisher CryoStar NX50 Cryostat) and 10 µm thick sections were cut. Multiple slices representing different z planes of the organoids were mounted on two warm, functionalized, bead-coated MERSCOPE slides. Slides were then placed in a 60 mm petri dish and stored in the cryostat for 20 minutes. Slides were dried at 50C for 30 min before washing the slides with 5mL 1X PBS (Invitrogen #AM9625) three times for 5 minutes each wash at room temperature to remove the OCT. Then, 5mL of 70% ethanol was added to the petri dish for permeabilization. Samples were photobleached for 3 hours to reduce the tissue background during imaging using the Vizgen Photobleacher (Vizgen #10100003). Samples were then washed with 5 ml Sample Preparation Wash Buffer (Vizgen, #20500001-2) before adding 5 ml Formamide Wash Buffer (Vizgen, #20500001-2) for 30 minutes at 37°C. The samples were then hybridized with the MERSCOPE Gene Panel Mix at 37°C in an incubator for 42 hours. The tissue slices were then washed twice with 5 ml Formamide Wash Buffer at 47°C for 30 minutes and embedded into a hydrogel using the Gel Embedding Premix (Vizgen, #20500001-2), ammonium persulfate (Sigma, A3678-25G), and TEMED (N,N,N’,N’-tetramethylethylenediamine) (Sigma, T7024-25ML) from the MERSCOPE Sample Prep Kit (Vizgen, #20500001-2). After 1.5 hours the gel embedding solution polymerized and the sample was cleared with a Clearing Solution consisting of 50 ul of Proteinase K (NEB, #P8107S) and 5 ml of Clearing Premix (Vizgen, #20500001-2) at 37°C overnight. Then, the samples were washed with 5 ml Sample Preparation Wash Buffer and imaged on the MERSCOPE system (Vizgen #10000001) using a MERSCOPE 500 gene imaging kit (Vizgen #10400012). A step-by-step instruction on the MERFISH sample prep is available at: https://vizgen.com/resources/fresh-and-fixed-frozen-tissue-sample-preparation/ and the instrumentation protocol is available at: https://vizgen.com/resources/merscope-instrument/.

#### Generation of doxycycline-inducible Cas9 nuclease stable cell line

The generation of a doxycycline-inducible Cas9 nuclease stem cell line was achieved through the stable integration of inducible Cas9 and reverse tetracycline-controlled transactivator (rtTA) into the AAVS1 safe-harbor locus using transcription activator-like effector nucleases (TALEN). To construct the donor vector containing Cas9 with dual nuclear localization signals (dual-NLS), the SV40-NLS and nucleoplasmin-NLS sequences were appended to the 5′ and 3′ ends of Cas9 (from the Puro-Cas9 donor, Addgene #392399) via PCR amplification. Prior to PCR, the Puro-Cas9 donor template was digested with AgeI and AscI to minimize background during cloning. The amplified dual-NLS Cas9 fragment was inserted into an AAVS1 Hygro-donor vector (Addgene #86883), downstream of a tetracycline-responsive element (TRE) promoter, using NEBuilder® HiFi DNA Assembly (NEB, E2621L) following the manufacturer’s instructions. The generation of the doxycycline-inducible Cas9 nuclease (iCas9) stem cell line was performed in accordance with previously published protocols^52,53^ with minor modifications^40^. Briefly, four plasmids, including dual-NLS-iCas9-Hygro, AAVS1-Neo-M2rtTA (Addgene #60843), AAVS1-TALEN-L (Addgene #59025), and AAVS1-TALEN-R (Addgene #59026), were prepared using endotoxin-free maxiprep kits (Qiagen, 12362) and nucleofected into WTC iPSCs. A total of 20 µg DNA with 8:8:1:1 ratio was nucleofected using the Lonza 4D-Nucleofector® X following the manufacturer’s protocol. Post-nucleofection, the WTC cells were plated on a 10-cm dish pre-coated with Matrigel in mTeSR+ medium supplemented with 5% CloneR (STEMCELL Technologies, #05888). G418 selection (100 µg/mL) was performed from days 2 to 5, followed by hygromycin selection (50 µg/mL) from days 7 to 9. Single clones were subsequently picked and screened for dual insertion of Cas9 and rtTA through genomic DNA PCR, followed by qPCR to assess doxycycline-inducible Cas9 expression^52,53^.

#### Cloning and lentivirus packaging for the perturbation experiment

Transcription factor knockout (TF KO) guide RNAs were designed utilizing CHOPCHOP and Synthego. To assemble vectors for each TF KO, gRNAs cassettes were assembled from PCR primers (IDT, Supplementary Table 5), amplified, and inserted into a template vector. Gel purified cassettes were assembled into a linearized vector via Gibson assembly (NEB, E2621). Presence of the correct dual gRNA cassettes were verified by Sanger sequencing. Lentivirus (LV) was produced in-house following a four plasmid second generation LV protocol requiring TAT-proteins for LTR dependent integration into the genome. A 1:1:1:1:4 plasmid mix consisting of 1.5μg of each Gag-pol, Vsvg, Rev and Tat helper plasmids with 6μg of the respective LV vector was used for transfection into 80-90% confluent HEK293T cells using TransIT-293 reagent (Mirus Bio, MIR 2705). Cells were cultured in high glucose DMEM (Gibco, 41965039) supplemented with 10% (v/v) FBS (Sigma-Aldrich, F2442), 1x Glutamax (Gibco, 35050061) and 0.2% (v/v) PS (Gibco, 15140122). To purify LV, the medium was changed to 30% FBS one day after transfection and collected after 24 hours. A 50x lentiviral suspension was generated in DPBS using Lenti-X™ concentrator (Takara Bio Inc., 631232). LV was produced separately for each knockout construct.

#### Stable cell line and mosaic organoids generation for perturbation experiment

A clonal doxycycline-inducible Cas9 iPSC line with WTC background (Coriell Institute), generated following previously established protocols^40^, was infected in a pooled fashion (Supplementary Table 5). As a control, LV harboring non-targeting guide sequences were used. Cells were cultured for 8 days after infection under continuous selection with 2 μg/ml blasticidin (Gibco, R21001). After selection and visual confirmation of fluorescent marker expression, cells were dissociated using TrypLE (Gibco, 12605010) and underwent further selection for positive fluorescence via FACS. Selected cells were plated and allowed to recover for 2-3 days (80-90% confluency) before being cryopreserved.

Cryopreserved cell pools were thawed and passaged once before cell suspensions were generated using TrypLE (Gibco, 12605010). A final cell suspension was created by mixing the KO pools (Supplementary Table 5) to ensure each KO was represented equally in the final cell suspension. The final mixed pool of cells was used to generate Qian organoids following the previously described method (Qian Reference). To induce Cas9 expression and therefore create knockouts in targeted genes, 2 μg/ml doxycycline (Clontech, 631311) was added with each media change starting on day 7 until day 14. Organoids were returned to their normal media on day 15. Fluorescence was monitored throughout culturing to ensure KOs reporters were not silenced throughout differentiation. Five organoids were pooled and dissociated into single cell suspensions at day 30 and day 70. Cells positive for fluorescent reporters were selected using FACS and these cells were then loaded onto the 10x Chromium using the Chromium Next GEM Single Cell 5’ v2 Dual Index kit (10x Genomics, PN-1000265) targeting 10,000 cells per reaction with two reactions per time point. Single-cell transcriptome and CRISPR gRNA libraries were prepared following manufacturer protocols. The transcriptome libraries and gRNA libraries were pooled and sequenced on the NovaSeq 6000 platform using 26/10/10/90 as cycle parameters.

#### Organoid culture for morphogen screen experiment

Brain organoids were generated from the WTC cell line. For the morphogen screen experiment protocol 1^13^ was used as a baseline and also serves as a control. Organoids were plated in ultra low-attachment plates (Corning, CLS7007), 5 organoids per condition. On day 15 they were transferred in 6-well plates, placing all 5 organoids per condition in one well and kept on shaker, as in the original protocol. The majority of morphogens were provided between day 8 and 14 (Supplementary Table 5), always using base medium as indicated in the original protocol. Exception being SHH, which was provided from day 1 to day 7, and CHIR, from day 5 to day 14, as in the original protocol; insulin, being provided from day 1 to 14 as in the published cerebellum protocol^18^; FGF-19 - from day 15 onwards, as it has been known to promote neural differentiation. We have used the following morphogens: SHH (R&D Systems, 1845-SH-025/CF), CHIR-99021 (Tocirs, 4423), insulin (Sigma, I9278), R-Spondin-3 (Peprotech, 120-44), R-Spondin-2 (Peprotech, 120-43), retinoic acid (Sigma-Aldrich, R2625), FGF-8 (STEMCELL Technologies, 78204), FGF-19 (Peprotech, 100-32), FGF-17 (Peprotech, 100-27), FGF-2 (Mitenyli Biotech, 130093840), BMP4 (Mitenyli Biotech, 130-111-167).

#### Single-nuclei isolation, fixation, snRNA-seq library preparation and sequencing for morphogen screen experiment

After 5 weeks (36 days) in culture 1-4 organoids per condition were dissociated individually using CyBio FeliX liquid handler robot with thermoshaker. For dissociation we used a papain-based dissociation kit as for the time course. Each organoid was dissociated using 820uL of enzyme mix 1 and 12uL of enzyme mix 2. Then each individual single cell suspension has been followed by nuclei isolation, as described for the time course (lysis time 2.5 minutes). Nuclei fixation and permeabilization procedures were performed according to the manufacturer specification (ParseBiosciences, nuclei fixation kit v2.1.2, WN400). Then collected samples were processed for highly multiplexed single-nucleus RNA sequencing (snRNA-seq) using a split-pool combinatorial barcoding kit (ParseBiosciences, WT Mega kit v2, dual-index version RX200).

### Data analysis methods

#### Preprocessing of single-cell multi-omic (scMultiome) data from the organoid time course

We used Cell Ranger ARC (v.2.0.0 with the default parameters to map the RNA-seq and ATAC-seq portion of the data to the human reference genome and gene annotation provided by 10x Genomics (GRCh38-2020-A-2.0.0). All samples were aggregated using the aggr command in Cell Ranger ARC to have the same list of accessible peaks being quantified for all cells. Both data modalities were read and further processed in R using Seurat (v.4.4.0)^54^ and Signac (v.1.1.1). As quality control, only cells satisfying the following criterias were remained: detected gene number between 1000 and 7500, percentage of mitochondrial transcripts less than 30%, detected ATAC-seq fragment number between 1000 and 30000, nucleosome signal less than 2.5 and TSS enrichment higher than 1.

Nuclei from different stem cell lines were demultiplexed using the demuxlet^55^ tool. Genotyping information was downloaded from the HipSci (WIBJ2, HOIK1) or Allen Institute (WTC) website. Bcftools were used to merge all vcf files and sites with the same genotypes in all samples were filtered out. Demuxlet was run with default settings on transcriptomic and genomic reads. Cells with ambiguous assignments were classified as “Unknown”. For all other cells, the best singlet assignment was considered.

To analyze the scRNA-seq modality, the standard log-normalization was firstly applied, and the top 3000 highly variable genes were identified. Subsequently, truncated principal component analysis (PCA) was performed with the scaled expression levels (across all cells) of highly variable genes as the input, using the RunPCA() function from the Seurat package. The first 20 principal components (PCs) were used to integrate different samples in the dataset (time point and protocol) using the CSS method^51^ (cluster_resolution=1.2). We performed UMAP^56^ to obtain a two-dimensional representation of data. We used the RunUMAP() function with default parameters using all components of the CSS matrix.

For the scATAC-seq modality, the tf-idf (term frequency times inverse document frequency) normalization, implemented as the default normalization method in the Signac package for scATAC-seq data, was firstly applied. Singular value decomposition (SVD) was then performed using the RunSVD() function from the Signac package, to the normalized counts of the top peaks. The first 50 latent semantic indexes, except for the first one for its high correlation with numbers of fragments per cell, were used for scATAC-seq data integration using CSS method (cluster_resolution=0.8), followed by performing UMAP for low-dimensional data representation.

#### Cell type annotation of the scMultiome data

To annotate the data, we firstly identified clusters using the RNA assay, based on the CSS integrated embeddings, with the FindNeighbors() (default parameters) and FindCluster() (resolution=0.5) methods implemented in Seurat. Clusters were annotated into cell classes (NPC, neuroblast, neurons, astrocytes, neural crest derivatives, mesenchymal cells) based on canonical marker expression. Neuron clusters were further annotated based on neurotransmitter transporter expression, into glutamatergic neurons, GABAergic neurons (not Purkinje cells), Purkinje cells, dopaminergic neurons and glycinergic neurons.

To further disentangle heterogeneity of dopaminergic neurons, we subsetted 4,171 cells that were classified as dopaminergic neurons and re-preprocessed scRNA-seq data: normalization, highly variable genes selection, scaling and integration. For integration with the CSS method first 20 PCs were used, with the parameter cluster_resolution=1.2. UMAP embeddings were obtained using the CSS matrix. Louvain clustering (resolution=0.5) was applied to identify subtypes of dopaminergic neurons. For dopaminergic neurons subtypes classification we used canonical markers from literature^57–60^. We have observed strong expressions of *FOXA2*, *LMX1A*, *NR4A2*, *EN1*, which indicate midbrain origin of dopaminergic neurons^61^. We detected subtypes that resemble ones previously described^58^ (hDA1), however we have not observed expression of *ALDH1A1* and *LMO3*, which would be specific for hDA2.

To further dissect the heterogeneity of glutamatergic neurons, we subsetted the 7,372 glutamatergic neurons we have annotated. For the scRNA-seq portion, 3000 highly variable genes were re-identified. Their scaled expression levels across cells were used to rerun PCA, with the first 20 PCs being used to run CSS integration across samples. To incorporate the scATAC-seq information, the weighted nearest neighbor graph approach, implemented as the FindMultiModalNeighbors() method in Seurat, was applied to estimate modal weights for each cells in the subset, resulting in a nearest neighbor graph incorporating both the scRNA-seq and scATAC-seq information. Louvain clustering (resolution=1) was then used to identify clusters, which were annotated based on marker expression and the mapping to the primary reference.

#### Mapping of scRNA-seq data to primary reference

We adapted the projection and comparison strategy as previously described^21^ to compare the transcriptomic portion of the scMultiome time course data to the human first trimester developing brain cell atlas^26^. In brief, the scANVI model and the primary atlas data were retrieved. The scRNA-seq portion of the time course data was projected to the primary scANVI latent space using scArches^27^. A bipartite weighted k-nearest neighbor (wkNN, k=50, weighting_scheme = “jaccard_square”) graph was constructed. Based on the reconstructed wkNN graph, cell class label transfer was done via a weighted majority voting. The regional labels were transferred to each cell in the time course data in a similar manner but hierarchically to take into account the brain region hierarchy, as described previously^21^.

Based on the reconstructed wkNN graph, for a given subset of the query data (e.g. all cells of one sample or cell type), a presence score can be computed for each cell in the reference atlas, as the sum of edge weights on the wkNN graph, as a metric of the likelihood that the cell type/state represented by the reference cell being present in the given query cell population. To summarize presence scores of multiple query populations, presence scores of each query population were min-max normalized across all the reference cells, followed by a max-pool across all query cell populations.

To estimate transcriptomic similarity between cells in the organoids and human developing brains, we adapted the matched metacell reconstruction strategy as described previously^21^. In brief, for each query cell, i.e. every cell in the time course data, the average expression of its neighbors in the reference was calculated, weighted by weights of the wkNN graph. Transcriptomic similarity was then quantified per cell as the Pearson correlation between a query cell and its matched metacell across highly variable genes.

#### Spatial transcriptomic data preprocessing and analysis

The raw images of DAPI and Poly-T were retrieved from the MERSCOPE data output folder. For each tissue section, organoid masks were created by binarizing the DAPI image (z=0), filling the holes, and identifying the connected pixels. For each organoid mask, the corresponding DAPI and Poly-T images were then cropped into its expanded bounding box (both x- and y-axis expanded for 500 pixels on each side).

For each cropped DAPI and Poly-T image, an intensity normalization was then applied, for each z- stack separately. In brief, for each image, the local minimum (denoted as x_0_) between the two peaks of its intensity distribution was firstly identified based on binning as one normalization anchor. The other normalization anchor (x_1_) was defined as the 99.9th percentile of the image intensities. The normalized intensity was then calculated as x’=(x-x_0_)/(x_1_-x_0_). Next, cellpose (2.0)^62^ was applied for segmentation, using the cyto2 model on the PolyT and DAPI channels. The segmentation was firstly done on each z-stack separately. The segmented masks were then stitched and harmonized across z-stacks using the stitch3D function in cellpose (Extended Data Fig. 3). Cell masks with fewer than 3,000 pixels were excluded. Furthermore, any scatter cell mask, defined as those with fewer than seven cell masks within 250 pixels of distance, was excluded.

Next, for each segmented cell after the filtering, transcripts being called by MERSCOPE analytic pipeline and overlapping with the cell mask were counted for each gene to quantify its gene expression profile. Cells with fewer than 40 detected transcripts were excluded. The transcript numbers per cell (*n*) were then normalized to the cell volumes (*v*, number of pixels) with a target sum of 40,000: *n*’=*n*/*v*\*40000.

To generate the UMAP of the MERSCOPE data, the volume-normalized expression levels of each of the 500 measured genes were log-transformed (with 1 pseudocount, the pp.log1p function in scanpy) and scaled (the pp.scale function in scanpy). PCA was applied to the scaled expression matrix. The 1st PC was discarded for its high correlation with numbers of detected transcripts per cell. The 2nd to 11th PCs were used to generate the neighbor graph using the pp.neighbors function in the scanpy package, followed by generating the UMAP embedding with the tl.umap function in the same package.

To transfer the cell type labels from the scMultiome data, we firstly generated a subset of the scMultiome data with matched time point and organoid culture protocol as the MERSCOPE data, by selecting cells from samples representing day-30 organoids generated with protocol 1. Cells with cell type labels of cell types with no more than 100 cells in the subset were further excluded from the subset. Next, we generated the UMAP embedding of the subset based on their CSS representation calculated earlier (with default parameters), as well as to do louvain clustering with a high resolution (resolution=5) based on its neighbor graph (with default parameters). For each of the high-resolution clusters, the average expression of the 500 genes measured in the MERSCOPE experiment was calculated, and the cell type label with the higher frequency among cells in the cluster was assigned to the cluster. Next, the expression profile of the 500 genes for each cell in the MERSCOPE data was correlated with each of the high-resolution clusters. The cell type label of the most correlated cluster was assigned to the MERSCOPE cell.

#### Characterization of chromatin accessible regions with cisTopics and differential accessibility analysis

We combined the cisTopic analysis and differential accessibility analysis to characterize the detected accessible regions in the scATAC-seq portion of the scMultiome data of the time-course atlas. To run cisTopic^28^ we used create_cistopic_object_from_fragments() in the pycisTopic package to generate a cell-count matrix from fragments. Considering the large size of the dataset, we run this analysis on the subsetted dataset (1000 cells per cell type). We performed topic modeling, varying the number of topics from 50 to 300 and selected the model with 250 topics. The estimated topic loadings were considered as the representations of the detected accessible regions. PCA was thus applied to the topic loadings, followed by louvain clustering (resolution=1) to identify region clusters based on the neighbor graph of accessible regions estimated with the first 50 PCs. To visualise clusters of the chromatin regions, a tSNE embedding was calculated (with default parameters). All the above analysis based on the cisTopic output was done with the scanpy package in Python (Supplementary Table 3, 4).

We also adapted a differential accessibility analysis^48^ to estimate statistical significance of accessibility differences among different cell types. In brief, the chromatin accessibility count matrix was firstly binarized. For each accessible region, two generalized linear models with quasibinomial error distribution were fit. The full model includes two independent variables: the number of detected fragments per cell, and the cell type label; the reduced model includes only the number of detected fragments per cell as the independent variable. The variation residuals of the two models were then compared using a F-test. The resulting p-values were adjusted using the bonferroni method. Accessible regions with the adjusted p-values<0.1 were considered as differentially accessible (DA) across cell types. Fisher’s exact tests were used to check the enrichment of DA regions in each region cluster.

#### Characterization of chromatin accessible regions clusters with GREAT and HOMER

We used rGREAT package^63^ and GO biological processes database to perform pathway enrichment analysis of the peaks clusters. For each cluster top 10 terms (based on lowest p-value adjusted using hypergeometric test over genes) were selected for presentation.

We used HOMER software^64^ and JASPAR 2024 database^65^, a subset of vertebrate motifs to perform motif enrichment analysis of peak clusters. For each cluster top 10 terms (based on lowest p-value) were selected for presentation. For both analyses only peaks, classified as differentially accessible (see previous section) were considered.

#### Mapping of scATAC-seq data to primary reference

First, we generated a cell-count matrix from fragments for organoid atlas using peak ranges, defined in the human developing brain chromatin accessibility cell atlas^30^ using the FeatureMatrix() function of Signac R package. Next, for each cell we inferred the topic distribution using peak counts information and topic-peaks loading of the cisTopic model presented in the primary cell atlas. Inference of topic profiles for query cells was done with an expectation-maximisation-like process. Each iteration includes: 1) computing the posterior probability of each topic given each peak by multiplying topic-peak loadings with the topic distribution vector from the previous iteration; 2) normalization across topics to make sure that probabilities are summed up to 1 for each peak; 3) refining the overall topic distribution using peak probabilities by multiplying the term frequency vector (peak counts vector) with the posterior probabilities of each topic given each peak; 4) re-normalisation of topic probabilities. For each cell, a maximum of 100 iterations were applied to estimate the topic probabilities. Alternatively, if the difference between an updated topic distribution and the previous one was less than 1e-4, the probability inference was considered to be converged, and the inference ended. The explained algorithm is adapted from lda python package^66^.

Next, to map the scATAC-seq data to the primary reference based on the inferred topic profiles, we applied scPoli to the primary scATAC-seq reference data, given the topic profiles (scaled across cells) as the input, with the donor information as the batch covariate, and the following information as the cell type labels: Cellclass and Celltype. The batch covariate was represented in the model as a learned vector of size five. We chose the hidden layer size of the one-layer scPoli encoder and decoder as 128, and the latent embedding dimension as 20. We trained the model for a total of 50 epochs, 35 of which were pre-training epochs. With the trained scPoli model, we used scArches to map the scATAC-seq portion of the scMultiome time-course atlas to the reference scPoli latent space. The inferred topic profiles were firstly scaled with the same scaling factors as the reference data. Next, the query model was fine-tuned with a batch size of 2,048 for a maximum of 20 epochs.

With the reference data and the query data represented with the same latent representations, we adapted the bipartite wkNN graph based comparison for label transfer, presence score estimation, and matched metacell reconstruction (on topic profile level) described above to compare the scATAC-seq portion of the scMultiome time-course organoid cell atlas and the human developing brain chromatin accessibility cell atlas.

#### Estimating cell line- and cell type-specificity for open chromatin peaks

In order to estimate connection between opened chromatin peaks and cell lines or cell types, we have utilized the following approach. Peaks counts in cells were binarised (if a peak had more than 1 count in the cell, the peak was considered as present). Then the mutual information between peak and assigned cell lines, conditioned on cell type, was calculated taking into account all cells of the dataset. The reverse procedure was also done, calculating mutual information between peak and cell types, conditioned on the cell line.

#### Calculation of motif enrichment scores for single-cell atlas

Transcription factor binding motifs (TFBSs), the sequence motifs that different transcription factors (TFs) recognize and bind, were retrieved from the JASPAR 2024 database^65^ using the JASPAR2024 R package in the format of position frequency matrices. The TFBS information was added to the scMultiome time-course atlas data with the AddMotifs() function in Signac. The chromVAR analysis^67^ was then run using the RunChromVAR() wrapper function in Signac (default parameters), to estimate the enrichment of each TFBS at each cell in the atlas.

#### GRN inference with Pando

First we have excluded non-neuronal cell types (mesenchyme and neural crest) and non CNS cell types (spinal cord neuroblasts). GRN initiation and motif search was done using default parameters of the Pando framework (v.1.0.0). GRN inference was done using highly variable genes and ATAC peak being aggregated to high resolution cluster level to reduce noise in data. We used a regularized linear model (“cv.glmnet”) with the elastic net mixing parameter 0.5. Modules were defined using find_modules_network() function with the following parameters: rsq_thresh = 0.1, nvar_thresh = 10, min_genes_per_module = 10. Module activity was calculated using Seurat function AddModuleScore() and taking in consideration target genes, inferred by Pando.

#### Estimating regional and cell type specificity of regulons for NPCs and neurons

To characterize the differences of TF usage in NPCs with midbrain and hindbrain identities, we extracted the subset of NPCs with midbrain and hindbrain regional annotation from the time-course data. Focusing on the 246 TFs which were detected in the data, with TFBS motif information in JASPAR 2024 and regulon inferred by Pando, Wilcoxon’s rank sum test, implemented as the wilcoxauc() function in the presto package in R, was applied to compare the gene expressions, corresponding TFBS enrichments (the chromVar estimated scores), and regulon activities between midbrain NPCs and hindbrain NPCs. TFs with significant differences (expression - adjusted P-value < 0.01, fold change > 1.2, AUC > 0.6, detection rate difference > 10%; chromVar score and regulon activity - adjusted P-value < 0.01, AUC > 0.7) were considered as showing regional differences in NPCs and potentially responsible for regional identity establishment.

To characterize the differences of TF usage in different neuron cell types, we firstly extracted the subset of neurons from the time-course data. To identify TFs representing different neuron cell types on expression, motif enrichment and regulon activity levels, we performed two different statistical tests on each level separately. Similar to the analysis on NPCs, we applied the Wilcoxon’s rank sum test implemented in the presto package as the first test to identify TFs with differences among cell types on different molecular levels (expression - adjusted P-value < 0.01, fold change > 1.2, AUC > 0.6, detection rate difference > 10%; chromVar score and regulon activity - adjusted P-value < 0.01, AUC > 0.7). In addition, we also implemented an F-test-based differential abundance test. The test compares the residuals of two linear models, the full model with both cell types and a neuron-NPC score as described previously^51^ as a covariate, and the reduced model with only the neuron-NPC score. The neuron-NPC score is calculated as differences on the module scores (with the AddModuleScores() function in Seurat) of neuron-high genes and those of NPC-high genes, and therefore represents the progress of neural differentiation. With the second test, any comparison with Benjemini-Hochberg adjusted p-value<0.05 was considered as statistically significant. For each of the molecular levels, we required a TF being significantly different among different neuron cell types if both tests suggest significant differences. Next, a TF was considered to contribute to the differences among different neuron cell types if its regulon shows differential activities, plus significant differences detected in either gene expression or motif enrichment level.

To compare regional differences of TF profiles in NPCs and neurons, we curated two lists of TFs with high-confidence of positive regulation to their targets. For NPCs, we required the differences between average levels in midbrain and hindbrain having the same sign for TF expression, the enrichment of their corresponding TFBS motifs, and their positive regulon activities. For neurons, we firstly calculated Pearson correlation of TF expression and positive regulon activity with motif enrichment for each TF across neuron cell types. The TFs with at least one correlation > 0.5 and neither <0.1 were considered to be high-confident positive regulators in neurons. The two lists were unioned, and their differences between their average levels in midbrain and hindbrain were calculated for NPCs and neurons separately. The profiles of the NPC and neuron regional differences were then correlated for each of TF expressions, TFBS motif enrichments and positive regulon activities.

To generate a low dimensional representation of TF specificity in different neuron cell types in the organoids, we focused on the high-confident positive TFs in neurons, and applied PCA to their scaled average regulon activities in different neuron cell types. The 1st and 5th PCs were selected as the 2D embeddings of TF neuron cell type specificity, with which TFs with the highest regulon activity at the same cell types tend to be separated. On the other hand, the cell type and regional specificity of TFs were also examined by comparing regulon activities and TFBS motif enrichment, with the assumption that TFs responsible for maintaining cell type identity of a neuron cell type should be high in both values.

#### Processing and analysis of CRIPSP-perturbation single-cell RNA-seq data

We used Cell Ranger (v.7.0.1) using flag *multi* to obtain count matrices of transcriptome and gRNAs. We used human transcriptome (hg38, provided by 10x Genomics), and a table of guide sequences for GEX and gRNA reads mapping respectively. To facilitate efficient gRNA detection, the pattern and sequence entries in the feature reference files were adjusted to 12bp instead of 20bp. Then, as with the time course, count matrices were further processed using the Seurat R package. Cells were filtered on the basis of the number of counts (>600,<20,000 for day 30) or detected genes (>500,<7500 for day 70) and the fraction of mitochondrial genes (<0.15 for day 30 and <0.1 for day 70). Transcript counts were normalized to the total number of counts for that cell, multiplied by a scaling factor of 10,000 and subsequently natural-log transformed (NormalizeData()). 10X lanes from the same time points were integrated using CSS method as for the time course data. Datasets were annotated combining label transfer from the time course dataset (using css_project() function of CSS package) and marker gene expression.

To assign gRNA labels, the 10x gRNA calling outputs (protospacer_calls_per_cell.csv) were first filtered to retain cells with gRNAs targeting a single gene. For cells with gRNAs targeting multiple genes, additional filtering was applied to identify the best call based on z-scaled UMI > 5 and a total UMI count > 10. Cells lacking gRNAs calls were excluded from further analysis. Cells with effective perturbations were identified using Mixscape^68^ as implemented in Seurat using cells with all other gRNAs as a control. Regulon activities upon CRISPR-based knockout were quantified using Seurat function AddModuleScore() considering the positive and negative regulons of TF of interest as inferred in Pando. To visualize the detected gRNAs and regulon activities on the UMAP embeddings, their densities were estimated using Nebulosa^69^ (weight=1 for gRNA). Differentially expressed genes (DEGs) were identified for each cluster with FindMarkers() function in the seurat package using wilcoxon test (log2 fold change >0.5, adjusted p<0.05, detection rate>5%, detection rate difference>10%) and analyzed for gene ontology enrichment using clusterProfiler^70^.Preprocessing of snRNA-seq data of morphogen perturbation screen

#### Processing and analysis of morphogen perturbation screen single-cell RNA-seq data

We used Parse Biosciences Software (v1.0.4) to demultiplex barcodes, map to hg38 human transcriptome and generate count matrix, which was further processed using scanpy python package (v1.10.3)^71^. Cells were filtered out on the basis of unique molecular identifier (UMI) counts (>1000, <30000) and the fraction of mitochondrial genes (<10). Then transcript counts were normalized to the total number of counts for that cell, multiplied by a scaling factor of 10,000 and subsequently natural-log transformed. Then highly variable genes were estimated and total UMI counts as well as fraction of mitochondrial genes were regressed out. After that PCA was performed, followed by neighbours estimation and leiden clustering.

An additional quality-control step included glycolysis signature calculation. We have followed the approach, described previously^21^, by selecting genes that belong to GO terms *’canonical_glycolysis’* and using tl.score_genes() function to estimate glycolysis score. The same procedure was done for *‘aerobic electron transport chain’*. Then, for each cell we calculated differences between glycolysis score and aerobic score and filtered out leiden clusters, which had median differences more than 0.05. After that, highly variable genes were calculated again, followed by scaling and PCA, as described above. Subsequently, we have calculated new clusters using leiden algorithm and UMAP embeddings to get 2D representation of data.

To annotate the data we firstly identified clusters using the RNA assay, based on the PCA embeddings, with the pp.neighbors(n_neighbors=30, n_pcs=60) and tl.leiden(default parameters) methods implemented in scanpy. Clusters were annotated into cell classes (NPC, neuroblast, neurons, neural crest derivatives) based on canonical marker expression. Neuron clusters were further annotated based on neurotransmitter transporter expression, into glutamatergic neurons, glycinergic neurons, motor neurons and dopaminergic neurons, and their subtypes were classified using canonical markers from literature^72–75,76,77^.

To further decipher heterogeneity of dopaminergic neurons, we have subsetted neuroblasts and dopaminergic neurons clusters (21,901 cells) and repeated reprocessing steps. To identify clusters we used pp.neighbors(n_neighbors=30, n_pcs=60) and tl.leiden(resolution=0.3) methods implemented in scanpy. Clusters were annotated into dopaminergic neurons subtypes based on the canonical markers from literature^57,59,78^. As in the time course atlas, we have not observed *CALB1+* or *ALDH1A1+* cells, and have *SOX6+, OTX2+* and *Vglut2+* subclusters, as well as A9-like subtype, expressing high level of *KCNJ6*.

#### Determination of composition changes in morphogen perturbation screen

To quantify differential cell type abundance under morphogen treatment in comparison to control, we have used speckle package^79^, which takes into account technical replicas of conditions. We used the propeller() function, using logit proportion transformation parameters. For each cell type, this function fits a linear model, and estimates p-value using t-test, subsequently performing FDR adjustment using Benjamini-Hochberg procedure. For estimation of differential dopaminergic cells abundance across conditions, we have considered only conditions that had more than 200 dopaminergic neurons cells, and followed the same steps as described above.

#### Estimation of anterior-posterior and dorsal-ventral scores

To train the models to estimate anterior-posterior (AP) and dorsal-ventral (DV) scores, the radial glia subset of a recently published first-trimester developing human brain transcriptomic atlas^26^ was used. In brief, the primary atlas was re-processed with scANVI^80^ with scVI^81^ model initialization plus cell type labels as the concatenation of dissection region and cell class. Details have been described previously^21^.

Next, radial glial cells (RGs) were selected for building the models. For the AP axis scoring model, the dissection tissues of the primary RGs were firstly summarized to telencephalon, diencephalon, mesencephalon and rhombencephalon. Cells with dissection tissue label being too broad (e.g. head and brain) were assigned to None. In order to correct for the potential label error due to mis-dissection, a label smoothening procedure was applied by calculating the numbers of neighboring cells dissected from each of the four summarized regions. A weighted proportion (weighted by Jacard index for neighborhood similarity) was calculated and the cell was reassigned to the region with the highest score. Based on the calibrated regional label, a value of 1, 2, 3 or 4 was assigned to each cell, 1 for telencephalon, 2 for diencephalon, 3 for mesencephalon and 4 for rhombencephalon. 5000 RGs with each of the four values were then subset to train a elastic net model (with glmnet package, default parameters) with the scANVI latent representation as the input.

For the DV axis scoring model, the intuitive dorsal and ventral scores were firstly calculated as the module activity scores of the dorsal organizer markers *(LMX1A, DMRT3, MSX1, BNC2, MSX2, EYA4, MAFB, RXRG, OLIG3*) and ventral organizer markers (*TOX2, NKX2-2, FOXA2, NKX6-1, FOXJ1, LMX1B, FOXA1, PHOX2B, SIM1, POU3F1,NKX2-1, SIX6, SIX3, RAX, MAF*) using the AddModuleScores function in Seurat. Both marker sets were derived from the mouse developing brain atlas^82^. The marker-based DV scores were calculated as the difference between the two modules scores (dorsal-ventral). A random subset of 80% of the RGs with absolute values of the marker-based DV scores > 5 were selected to train an elastic net model (with glmnet package, binomial family) to classify dorsal and ventral cells with the scANVI latent representation as the input.

To apply the two scoring model to the organoid data, scArches^27^ was firstly used to project the organoid data to the scANVI model of the primary atlas to obtain the projected latent representation. Next, the projected representation was used as the input to the trained AP and DV axis scoring model for prediction (type=‘link’).

#### Mapping to primary data and estimation of gained populations

To map morphogen screen data to primary human developing brain atlas we utilized the approach described previously^21^ and above. Briefly, scArches were used to map scRNA-seq data to the scANVI model of the human developing brain atlas. Sublibrary variable was chosen as a batch covariate. We trained the model for 100 epochs with weight_decay = 0 and otherwise default parameters. After mapping we have calculated max presence scores of each condition for primary cells.

#### Trajectory inference based on cluster co-occurrence

We obtained the cell type proportions for different morphogen treatment conditions, and calculated the pairwise Pearson correlation (*r*) between cell types based on their proportions in different conditions. Using 1-*r* as the distance, the cell type dendrogram and hierarchical clustering with the complete linkage function was applied to cell types annotated in the morphogen screening data. The number of clusters (k=9) was determined manually to separate as many as possible different NPC populations and avoid neuron-only clusters. The resulting clusters of cell types were annotated based on the regional identities of the involved cell types.

#### Morpho-GRN inference with SCENIC

To infer the regulatory mechanisms of how different morphogens regulate changes on neural cell fate decisions, we adapted the morphogen-TF-regulon regulatory network inference approach described previously^40^. In brief, we firstly applied SCENIC to the morphogen screening scRNA-seq data to infer TF-target regulationship. We generated 20 random subsets of the data, in each of which 400 cells per cell cluster (excluding cells in the neural crest and melanocyte clusters) were randomly selected. GRNBoost2 was then applied to each of the subset with the arboreto_with_multiprocessing.py code in pySCENIC (all with a given seed of 10). The TF-target pairs appeared in at least three GRNBoost2 outputs were kept, with the max weights among all the estimated weights being used as the summary. The motif database v9 from cisTarget was then loaded to do trimming based on TFBS motif enrichment based and correlation, given the summarized GRNBoost2 results and each of the 20 random subsets. Only TF-target pairs appearing at least three times among the 20 trimmings were kept as the final results. For each TF, the union of its predicted targets is considered as its regulon, and regulons with fewer than five genes were discarded. Next, the regulon activities were calculated with the tl.score_genes() function in the scanpy package in Python.

To then infer the regulatory relationship between morphogens and TF regulons, we ran GRNBoost2 with default parameter and seed=10 using the arboreto_with_multiprocessing.py code in pySCENIC, given morphogen treatments (concentrations as values) as putative regulators and TF regulons (regulon module scores as values) as the putative targets. This analysis was only applied to the NPCs of the data, assuming that morphogen effects were on the commitment of different NPC populations.

Since morphogen usages are unbalanced across conditions, we designed a subsampling strategy to target different numbers of cells for different conditions (Extended Data Fig. 9). Firstly, we performed a hierarchical clustering of conditions based on their cell type enrichment relative to the control conditions, generating eight condition clusters. For each of them, conditions were further grouped into morphogen usage groups, in each of which conditions use the same morphogen combinations regardless of concentrations. Next, we defined a cap of 1,000 NPCs as the targeted NPC number for every condition cluster during subsampling. This targeted number of NPCs per condition cluster was evenly assigned to its morphogen usage groups. Lastly, the targeted number of NPCs per morphogen usage group was again evenly assigned to the conditions, generating the target NPC number for each condition. During each subsampling, if a condition includes more NPCs than the targeted number, the target number of NPCs was randomly selected; otherwise, all NPCs of the condition were always included. This random subsampling procedure was repeated 50 times to generate 50 NPC subsets.

Given each of the NPC subsets, GRNBoost2 was applied to estimate regulatory strengths of morphogens on each TF regulon. For each morphogen-regulon pair, the estimated importances with the 50 different NPC subsets were summarized using maximum. A threshold of estimated importance > 100 was used to screen for confident morphogen-regulon regulations. In addition, Pearson correlation (*r*) was estimated between each morphogen usage concentration and regulon activities across all NPCs, and any morphogen-regulon pair with *r*<0.1 were further discarded.

#### Trajectory inference based on transcriptome similarity

Based on the mapping of the morphogen screening scRNA-seq data to the primary human developing brain atlas, as described above, we calculated the max-min normalized presence scores of each cell type annotated in the screening data for all cells in the primary atlas, similar to what has been described above. For each cell in the primary atlas, the maximum across presence scores of all cell types in the screening data (cell-type-max presence scores) was calculated. Two criterias were used to identify primary cells matched with the screening cell types: 1) cell-type-max presence scores > 0.9; 2) cells in the primary cell cluster in which more than 80% of cells show cell-type-max presence scores > 0.5. Any primary cell satisfying at least one of the two criterias was considered as a matched primary cell, and was included for the primary trajectory analysis (the matched primary sub-atlas).

Next, we subsetted the matched primary sub-atlas and re-identified 5,000 highly variable genes using the pp.highly_variable_genes() function in scanpy (batch_key=’Donor’). Next, PCA was applied to the scaled expression of the identified highly variable genes, followed by harmony integration^83^ to align the top-20 PCs of different donors. The UMAP embeddings for visualization of the primary sub-atlas was generated based on the harmony integration result.

To identify the matched population to the ventral medulla trajectory in the screening data, two cell-type-max presence scores were calculated for each cell in the matched primary sub-atlas: the max presence scores for the ventral medulla cell types (*s_VM_*) including glutamatergic neuron (RELN high, glycinergic neuron (PAX2 high), glycinergic neuron (POU6F2 high, neuroblast Medulla, and NPC V-Medulla; and the max presence scores of other cell types (*s_O_*). Cells with *s_VM_*-*s_O_* > 0.1 were considered as the ventral medulla matched primary population.

To compare the matched ventral medulla trajectory in the two systems, we calculated NPC and neuron module scores with the AddModuleScore() function in Seurat for the screening data and the matched primary sub-atlas, given the NPC and neuron markers as described above. Next, we focused on the ventral medulla trajectory in the screening data, and the matched ventral medulla trajectory in the primary sub-atlas. Genes were tested in each of the two data for their expression in relation with the neuron-NPC scores representing neural differentiation progress. This was done by using a F-test-based approach as described in a previous study^48^, to compare the residuals of two linear models: the full model includes natural splines on the neuron-NPC scores (df=5), and the reduced model with only the intercept. Genes with Benjemini-Hochberg corrected p-value<0.01 were considered as differentiation-related differentially expressed (DE). For the union of genes with differentiation-related DE in the two systems, we used a cubic smoothing spline model, implemented by the smooth.spline() function in R, to evenly interpolate expression along the trajectory (with neuron-NPC scores in between -0.5 and 1.2). For each gene, Pearson correlation was calculated between the interpolated expression in the screening data and the primary data.

To validate the trajectories identified in the morphogen screening data based on cluster co-occurrence, we firstly defined the confidently matched primary neuron populations to different neuron cell types identified in the morphogen screening data. In brief, we focused on primary cells with the neuron-NPC score > 0.75, and assigned a cell to a neuron cell type if its normalized presence score is at least 0.5 higher than the normalized presence score of any other neuron cell types. Similarly, we defined the confidently matched primary NPC populations to different NPC cell types identified in the morphogen screening data, focusing on primary cells with neuron-NPC scores < 0.

With both the neurons and NPCs labeled in the matched primary-subatlas, we performed two different trajectory inference analysis on the matched primary sub-atlas. The first approach is CellRank2 with a hybrid of pseudotime kernel (with neuron-NPC scores as pseudotime, 50%) and connectivity kernel (with the unintegrated PCA, 50%), given neurons confidently matched to different neuron cell types as terminal states. The estimated likelihoods to different neuron cell types were summarized for NPCs confidently matched with different NPC populations in the screening data. The second approach is a stepwise neuron cell type label prediction along the differentiation trajectory, principally similar to FateID^84^. In brief, we binned cells in the matched primary sub-atlas into four groups based on their neuron-NPC scores: (-∞, -0.5], (-0.5, 0.25], (0.25, 0.75], (0.75, ∞). Starting from the last group, we trained an elastic net model with multinomial family using the glmnet package in R giving the assigned matched neuron cell type labels as the response variable and the harmony integrated PCs as the independent variables. The trained model was then applied to generate label prediction for the next group, and a new model was trained using cells in the next group given the predicted label. This procedure was applied iteratively until the neuron cell type labels were assigned to all cells. Lastly, frequencies of predicted neuron cell type labels were counted for NPCs assigned to different NPC populations.

## Acknowledgments

We thank all members of the Treutlein and Camp labs for discussions and feedback. We thank Sten Linnarsson for access to the human developing midbrain scRNA-seq data prior to publication; Camiel C. A. Mannens for sharing the cisTopic analysis results of the human developing brain scATAC-seq atlas; Jasper Janssens for discussion on the design of cerebellum conditions in the morphogen screen and morpho-GRN analysis; Aline Xavier da Silveira dos Santos, Benedikt Eisinger, and Fides Zenk for supporting generation of the doxycycline-inducible Cas9-nuclease iPS cell line; Fátima Sanchís-Calleja for insightful discussions on morphogens, patterning and brain regionalization; Jonas S. Fleck for insightful discussion on GRN inference, regulons and high resolution clustering. We thank the Genomics Facility Basel for sequencing, the ETH Zurich D-BSSE Single Cell Facility for support with cell sorting and imaging.

H.-C.L. was supported by a Human Frontier Science Program fellowship (LT000399/2020-L), and an EMBO long term fellowship (ALTF 1190-2019). The work was co-funded by the European Research Council (Braintime-874606, B.T.), the Swiss National Science Foundation (Project Grant 310030_192604, B.T.), the National Center of Competence in Research Molecular Systems Engineering (B.T.) and the Roche Institute of Human Biology (N.A., Z.H., H.-C.L., B.T.).

## Authors contribution

H.-C.L. performed organoid culture for the time course experiment, with support from M.S. and advice from S.K.; H.-C.L. generated scMultiome time course dataset with support from M.S.; B.K. performed the CRISPR-perturbation experiments with the support from A.M.; N.A. and B.K. generated and cultured organoids, used for morphogen screen; N.A. generated single-cell transcriptomic dataset of morphogen screen with support from H.-C.L., M.S, R.O. and V.B.; M.N. generated IHC data; R.H. established automatic organoid dissociation with support from N.A.; A.M. generated spatial transcriptomic data.

N.A. and Z.H. performed the analysis of scMultiome time course data with support from H.-C.L.; T.T. and Z.H. performed the analysis of spatial transcriptomic data; H.-C.L. performed the analysis of CRISPR-perturbation data with support from Z.H. and N.A.; N.A. performed the analysis of morphogen screen data with support from Z.H. and H.-C.L.; Z.H. performed morphoGRN and trajectory analysis of morphogen screen. N.A., Z.H., H.-C.L, J.G.C. and B.T. designed the study and wrote the manuscript.

## Data and code availability

The count matrices and metadata of the RNA-seq portion of the time course single-cell multiomic data is part of the previously published integrated human neural organoid cell atlas which is available at zenodo (https://doi.org/10.5281/zenodo.11203684) and the CellxGene Discover Census (https://cellxgene.cziscience.com/collections/de379e5f-52d0-498c-9801-0f850823c847). The raw and processed data of the scATAC-seq portion of the single-cell multiomic data as well as the scRNA-seq data of the morphogen screen will be uploaded to Array Express.

All code generated in the study, including analysis parameters, will be made available at https://github.com/quadbio.

## Extended Data Figures

**Extended Data Fig. 1.**
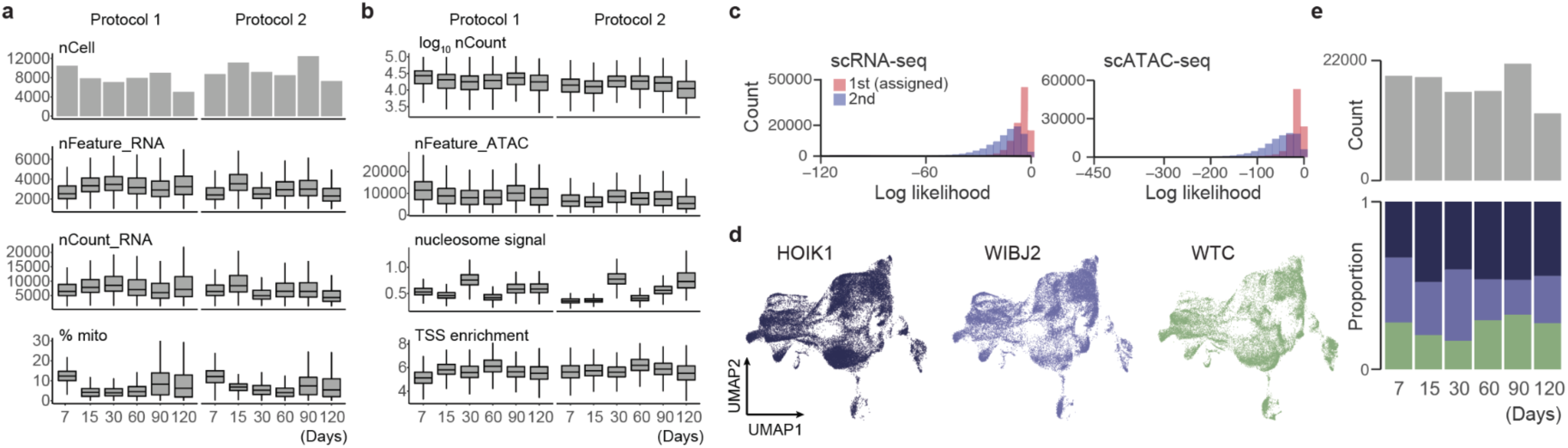
Quality control analysis of time-course single-cell multi-omic data. (a, b) Quality control of the time-course snRNA-seq (a) and snATAC-seq (b) in 2 protocols. Each column represents a time point. nCell, number of cells; nCount_RNA, total number of RNA molecules detected within a cell; nFeature_RNA, total number of genes detected within a cell; %mito, percentage of mitochondrial genes; nCount, number of detected reads; nFeature_ATAC, number of peak region fragments; TSS, transcription start site. (c) Histograms showing assignment log likelihoods for demultiplexing based on single nucleotide variants. (d) iPS cell (iPSC) lines distribution on the UMAP embedding. (e) Bar plot showing number of cells (top) and stacked barplot showing proportion of cell lines (bottom) at different time points.

**Extended Data Fig. 2.**
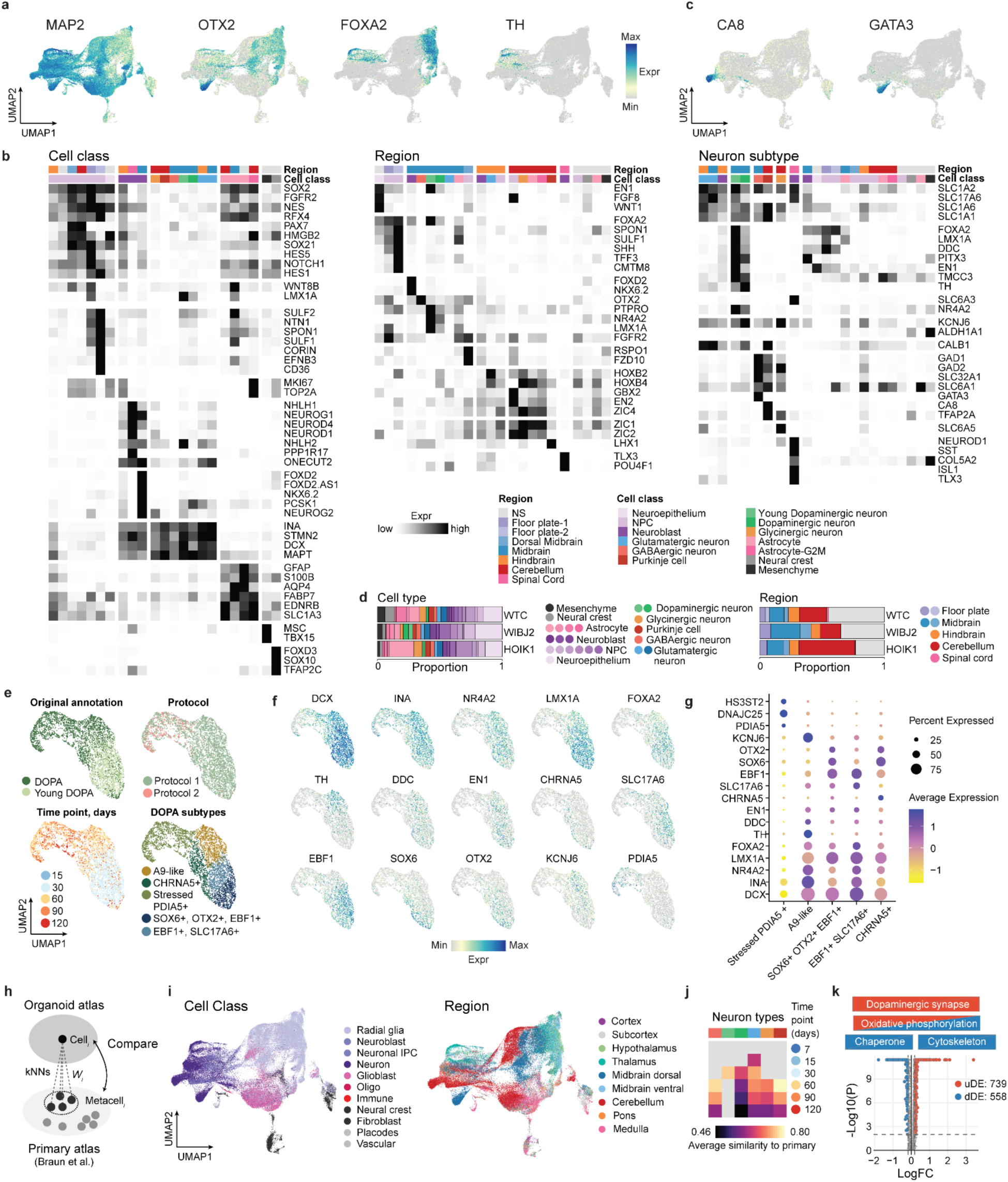
Supplemental analysis of time-course single-cell transcriptomic data. (a) UMAP embedding colored by marker gene expression, used for immunohistochemical stainings (log(transcript counts per 10k+1)). (b) Heatmaps showing min-max scaled mean expression of marker genes for each cell type. Genes are grouped by cell class markers (left), region markers (center), neuron subtypes markers (right). (c) UMAP embedding colored by marker gene expression (log(transcript counts per 10k+1)). (d) Stacked barplots showing proportions of cell types (left) and brain regions (right) for different iPSC lines. (e) UMAP embedding of dopaminergic neurons subset, colored by original cell type annotation, protocol, time point, dopaminergic neurons subtypes. (f) UMAP embedding of dopaminergic neurons, colored by marker gene expression (log(transcript counts per 10k+1)). (g) Dotplot showing marker genes expression in dopaminergic neurons subtypes. (h) Schematic of atlas projection to the primary human developing brain scRNA-seq cell atlas. (i) UMAP embedding colored by the scArches-transferred cell class (left) and region (right) labels from the primary reference. (j) Heatmap showing average similarity of neuronal subtypes to primary counterparts. (k) Volcano plot showing differentially expressed genes between brain organoid atlas dopaminergic neurons and primary dopaminergic neurons.

**Extended Data Fig. 3.**
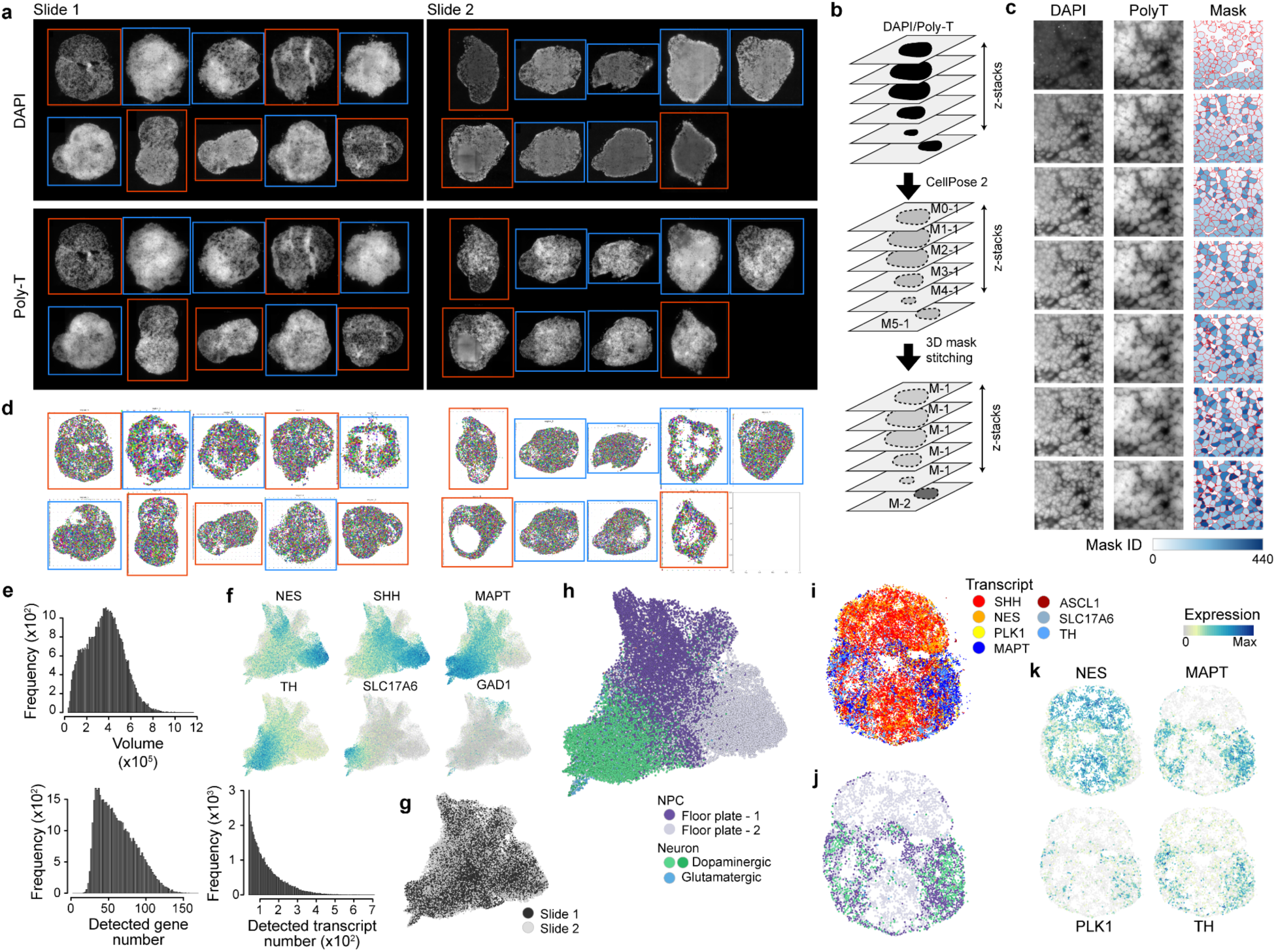
Supplementary analysis of the spatial transcriptomics data. (a) DAPI and polyT signal for each slide. Orange and blue frames indicate tissue sections from the two different organoids (b) Schematic illustrating MERFISH data analysis pipeline. (c) Example images, showing crop section from spatial transcriptomic slide. Left column shows DAPI signal, middle - polyT signal, right - inferred cell masks. (d) Slices, showing overlapped MERFISH probe signals. (e) Histograms, showing distribution of segmented cells volume (top), detected gene per cell (bottom left), detected transcript per cell (bottom right). (f) UMAP embedding of segmented cells, based on their summarized transcriptomic features, colored by marker gene expression. (g) UMAP embedding colored by slide. (h) UMAP embedding colored by cell type label, transferred from corresponding scRNA-seq data. (i-k) Example of organoid slice profiled at slide 2, showing selected transcript detection (i), transferred cell type label (j), marker gene expression (log(transcript counts per 10k+1)) (k).

**Extended Data Fig. 4.**
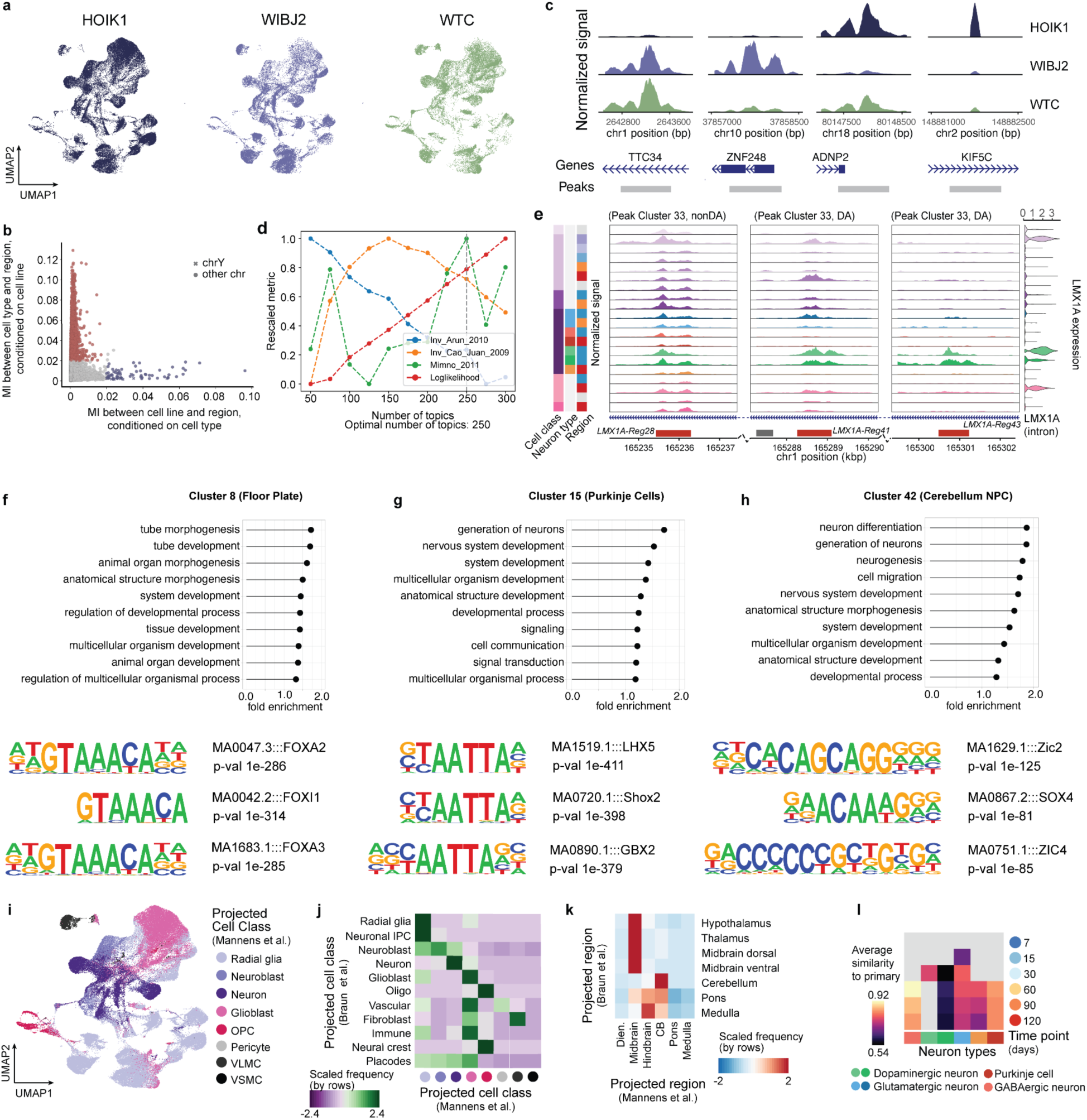
Supplemental analysis of time course single-cell chromatin accessibility data. (a) iPS cell (iPSC) lines distribution on the UMAP embedding of sc chromatin accessibility data. (b) Scatterplot, visualising mutual information between cell line and open chromatin region, and cell type and open chromatin region for each peak. Shape of the point indicates whether the peak is mapped to the Y chromosome. (c) Chromatin accessibility profile tracks in different cell types at 4 representative loci, shown to be cell line-specific. (d) Visualisation of metrics value for models with different numbers of topics. All metrics are min-max scaled, Arun and Cao metrics were inverted for visualisation purposes. (e) Chromatin accessibility profile tracks in different cell types at three representative loci for *LMX1A* gene. Violin plots on the right show *LMX1A* split by cell type. (f-h) Cluster characterisation. Examples of enriched GO-biological processes terms from GREAT analysis of DA peaks of the indicated cluster (top) and enriched motifs (bottom) for cluster 8 (f), 15 (g) and 42 (h). All adjusted p-values of presented terms were < 10^-30, hypergeometric test over genes. Fold enrichment is also derived from hypergeometric test over genes. (i) UMAP embedding colored by the transferred cell class from the primary reference. (j-k) Heatmap showing consistency between projected labels from the transcriptomic (rows) or chromatin accessibility (columns) primary atlases, for the cell class labels (j) and region/tissue labels (k). (l) Heatmap showing average similarity of neuronal subtypes to primary counterparts, based on opened chromatin profiles.

**Extended Data Fig. 5.**
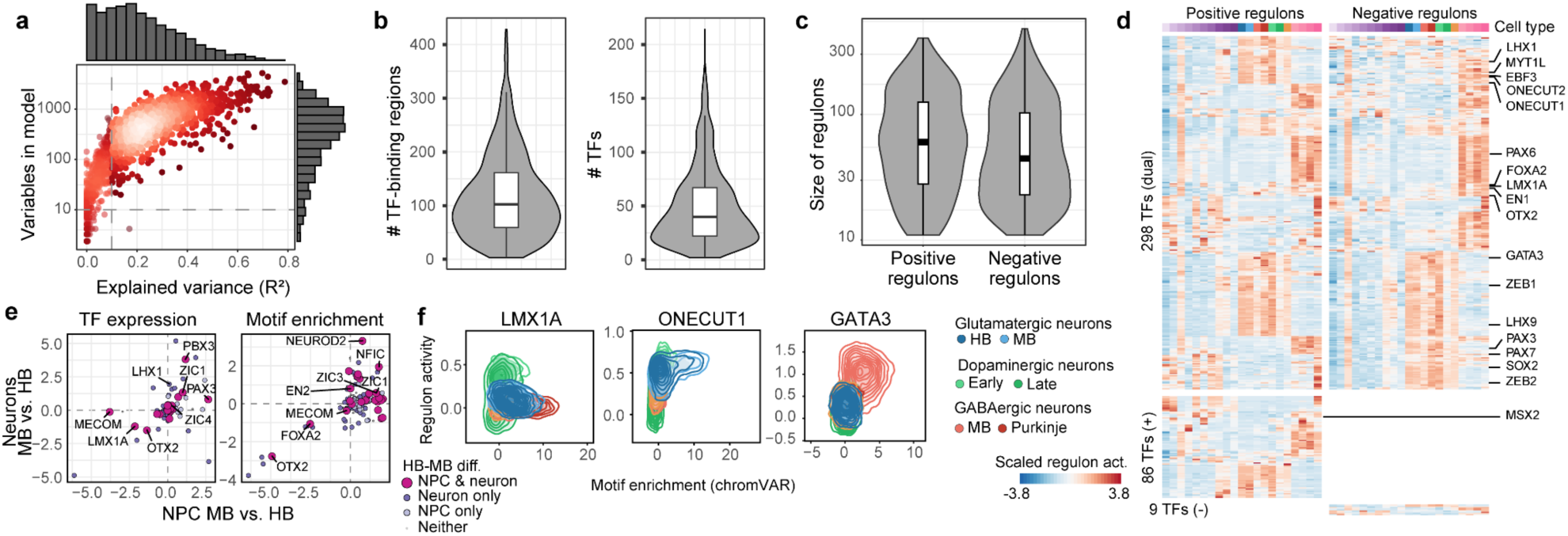
Supplementary analysis of the regulon inference using the PANDO framework. (a) Scatter plot and histograms indicating explained variance versus number of variables of models for GRN construction. (b) Violin plots showing the distribution of numbers of predicted binding regions (left) and TFs (right) per target gene in the GRN. (c) Violin plots showing size of regulons (number of genes per TF), split by positive and negative regulons. (d) Positive and negative regulon activity, averaged by cell type. 298 TFS showed dual activity (acting as repressors for some genes and activators for others), 86 TFs showed only positive activity, and 9 TFs only negative. (e) Comparison of regional differences of the TFs expression (left) and motif enrichment in open chromatin (right) for neurons (y-axis) and for NPC (x-axis). Values were summed up based on regional identity and then the difference between hindbrain and midbrain was calculated for each TF. The size and color of the dots represent the results of the DE analysis of the regulon activity between midbrain and hindbrain: if the regulon was in DE for both neuron and NPC, only in one of them, or in neither. MB, midbrain; HB, hindbrain. (f) Density plot, indicating motif enrichment and regulon activity for selected transcription factors for each neuron subtype.

**Extended Data Fig. 6.**
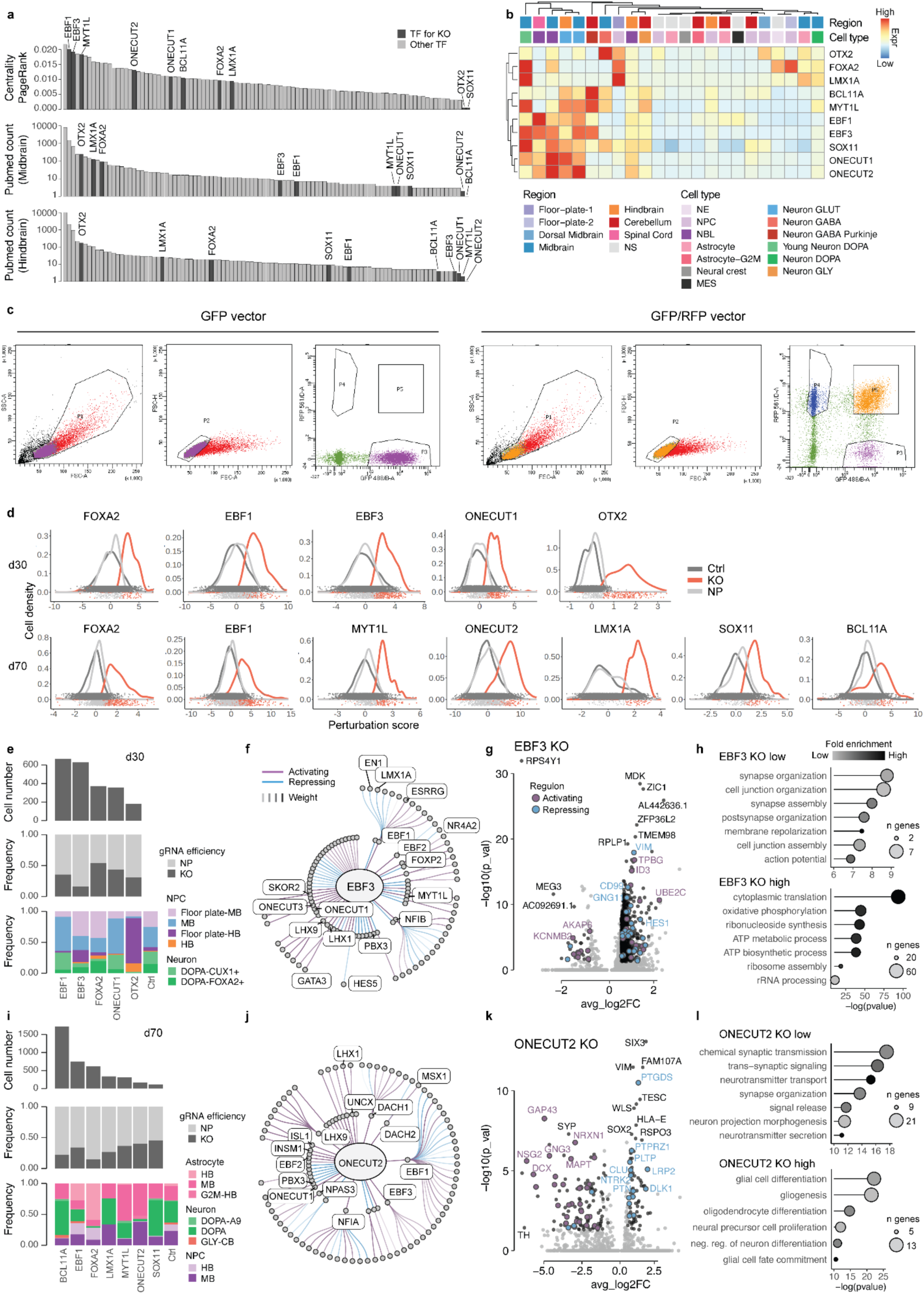
Selection of target genes for single-cell perturbation experiment and supplementary analysis of CRISPR-screen data. (a) Barplots showing PageRank regulon centrality (top), number of papers, in which terms TF name and “Midbrain” co occur (middle) and number of papers, in which terms TF name and “Hindbrain” co occur (bottom) for each TF. (b) Heatmap showing average expression of TFs, chosen for KO experiment, split by cell type. GLUT, glutamatergic; GABA, gabaergic; DOPA, dopaminergic; GLY, glycinergic; NS, non specific. (c) Exemplary Fluorescence-activated cell sorting plots of the sorting scheme used to isolate GFP vector positive (left) and GFP/RFP vector positive(right) iPS cells. (d) Distributions of perturbations scores for TF-targeting gRNAs, split by guide efficacy classification. Ctrl, guide was not detected; NP, non perturbed: guide is detected, but not efficient; KO, knock-out. (e, i) Barplots, showing number of cells, carrying efficient guide (top), proportion of efficient guides (middle), proportions of cell types (bottom) for each target KO gene for day 30 time point (e) and day 70 (i). MB, midbrain; HB, hindbrain; DOPA, dopaminergic neurons, NP, not perturbed; KO, knock-out. (f, j) GRN subgraph for *EBF3* (f) and *ONECUT2* (j), showing first- and second-order TF targets (nodes). The edges are colored based on TF regulatory interaction: purple is activating, blue - repressing. (g, k). Volcano plot showing DE between *EBF3*-KO and control cells for day 30 time point (g) and between *ONECUT2*-KO and control cells for day 70 time point (k). DEGs are colored in black. DEGs belonging to positive and negative regulons are further labeled in purple and blue, respectively. (h,l) GO analysis for DE gene sets identified in g and k. Rows represent GO term, x-axis represent p- values and size of the dots is proportional to the gene set size.

**Extended Data Fig. 7.**
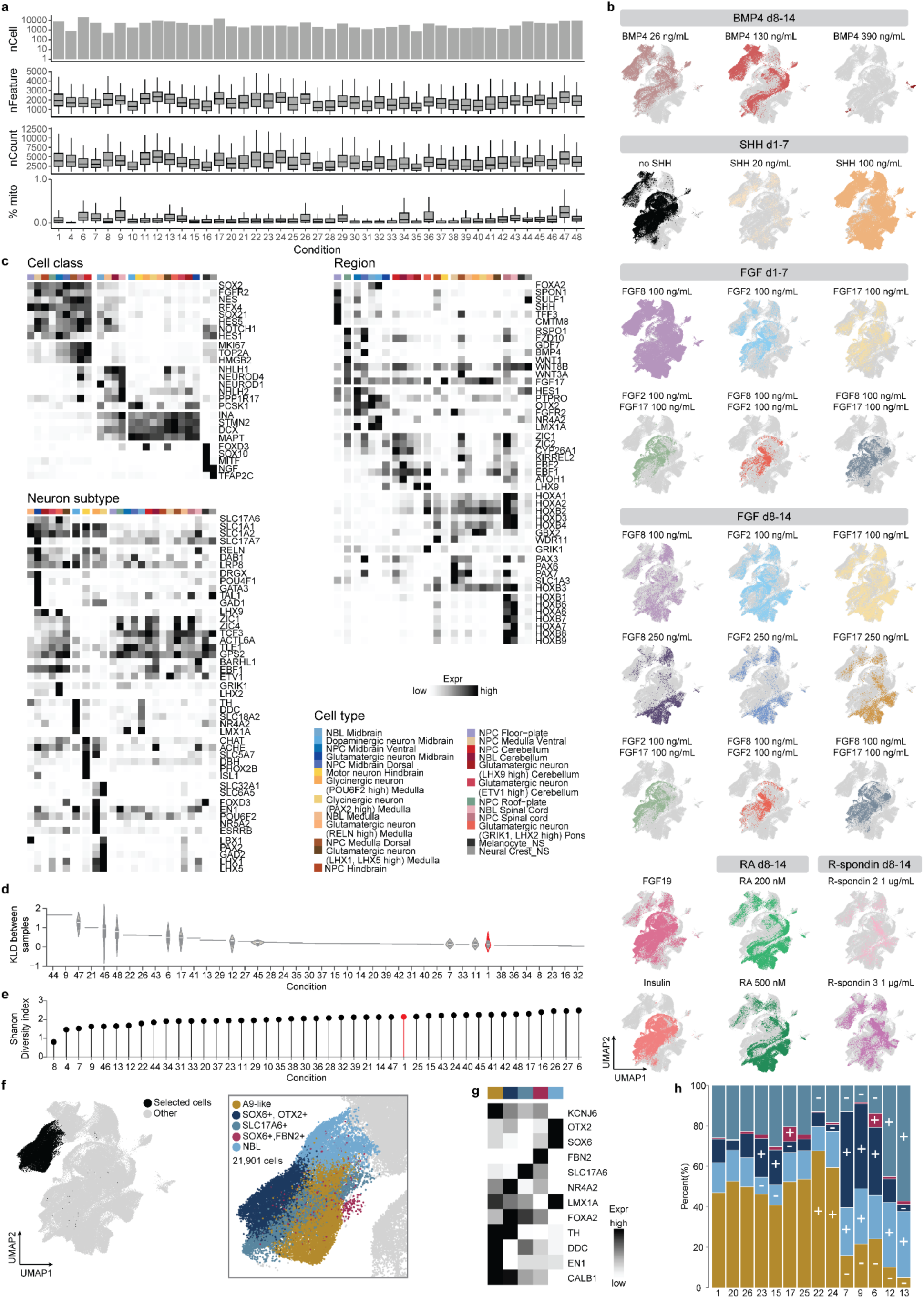
Supplementary analysis of morphogen screening experiment. (a) Quality control of the morphogen screen snRNA-seq. Each column represents a treatment condition. nCell, number of cells; nCount_RNA, total number of RNA molecules detected within a cell; nFeature_RNA, total number of genes detected within a cell; %mito, percentage of mitochondrial genes; nCount, number of detected reads. (b) UMAP embeddings, coloured by provided morphogen. For each panel treated cells are shown in colour, all other cells - in gray. (c) Heatmaps showing min-max scaled mean expression of marker genes for each cell type. Genes are grouped by cell class markers (top left), region markers (right), neuron subtypes markers (bottom left). (d) Violin plots, showing pairwise Kullback–Leibler divergence measured between organoids replicas. Control condition is highlighted in red. KLD, Kullback–Leibler divergence. (e) Lolipop plot, showing Shannon diversity index for each condition, calculated based on cell type proportions. Control condition is highlighted in red. (f) UMAP embedding, cells, which were selected for the subsequent dopaminergic neurons analysis (left) and cropped panel from the same UMAP representation, colored by dopaminergic neurons subtypes. (g) Heatmaps showing min-max scaled mean expression of marker genes for each cell type. (h) Stacked barplot, showing distribution of dopaminergic neuron subtypes in conditions, where more than 300 of these cells were detected. + and - indicate cell types, which were significantly enriched or depleted in this condition. p-value <0.05 (FDR adjusted using Benjamini-Hochberg procedure).

**Extended Data Fig. 8.**
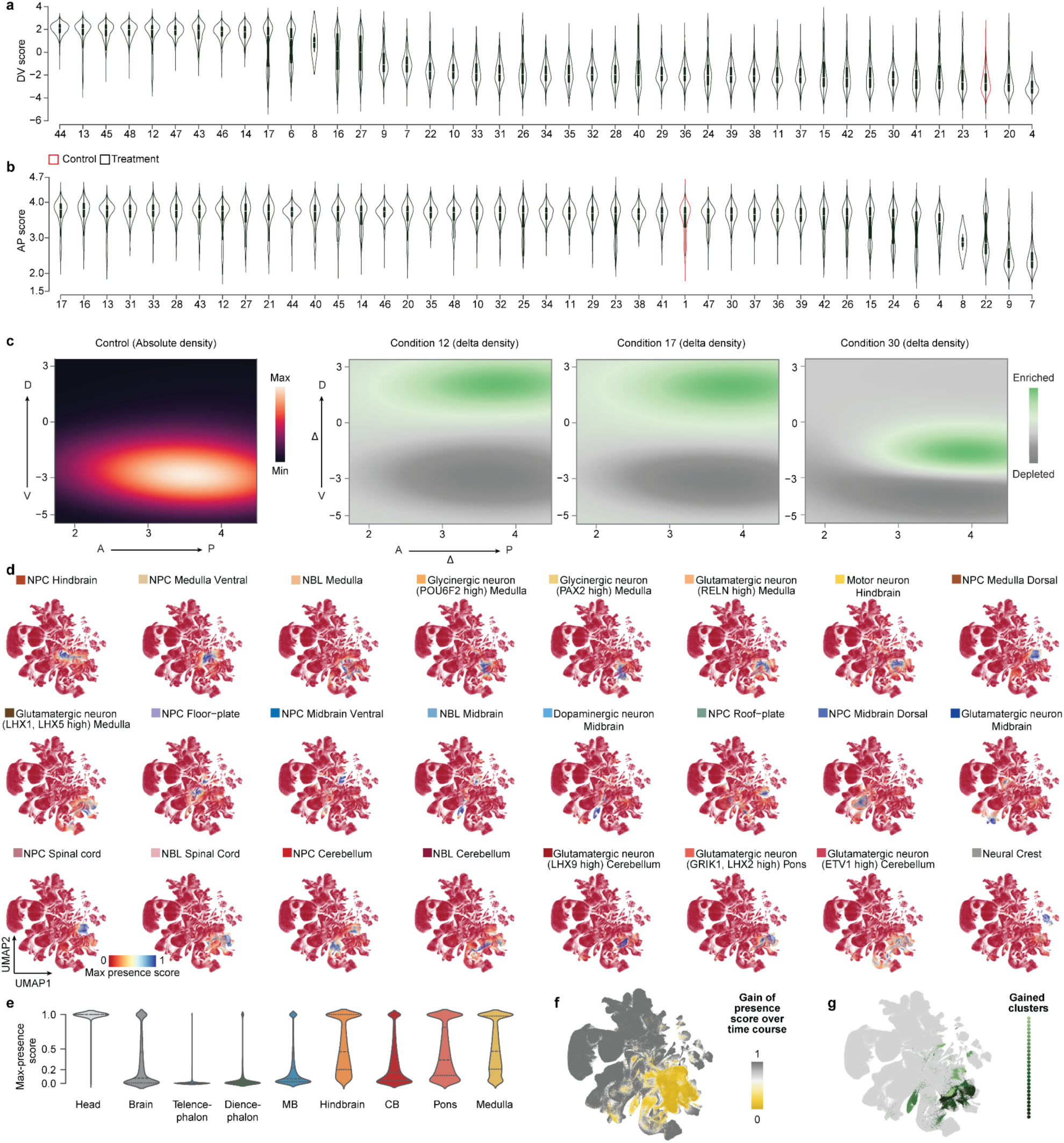
Supplementary analysis of mapping morphogen screen experiment to primary human brain atlas. (a, b) Violin plots, showing distribution of dorsal-ventral (a) and anterior-posterior (b) scores for each condition, ordered by descending median. Control condition is shown in red. DV, dorsal-ventral; AP, anterior-posterior. (c) Density plot, showing distribution of AP and DV scores for each cell in the control condition (left). Visualisation of difference of spatial mapping of selected morphogen treatment conditions in comparison to control using the dorso-ventral (vertical) and anterior-posterior (horizontal) axis of the developing human brain radial glia^26^. D, dorsal; V, ventral; A, anterior; P, posterior. (d) UMAP of the primary reference, colored by the max presence scores across the indicated cell types, detected in morphogen screen data. (e) Violin plots, showing distribution of max presence scores of morphogen screen data in primary data , split by dissected regions of primary data. (f) UMAP of the primary reference, colored by gained coverage of posterior brain organoid morphogen screen over posterior brain organoid time course atlas with negative values trimmed to zero. (g) UMAP of the primary reference, colored by gained cell types clusters in the morphogen screen data.

**Extended Data Fig. 9.**
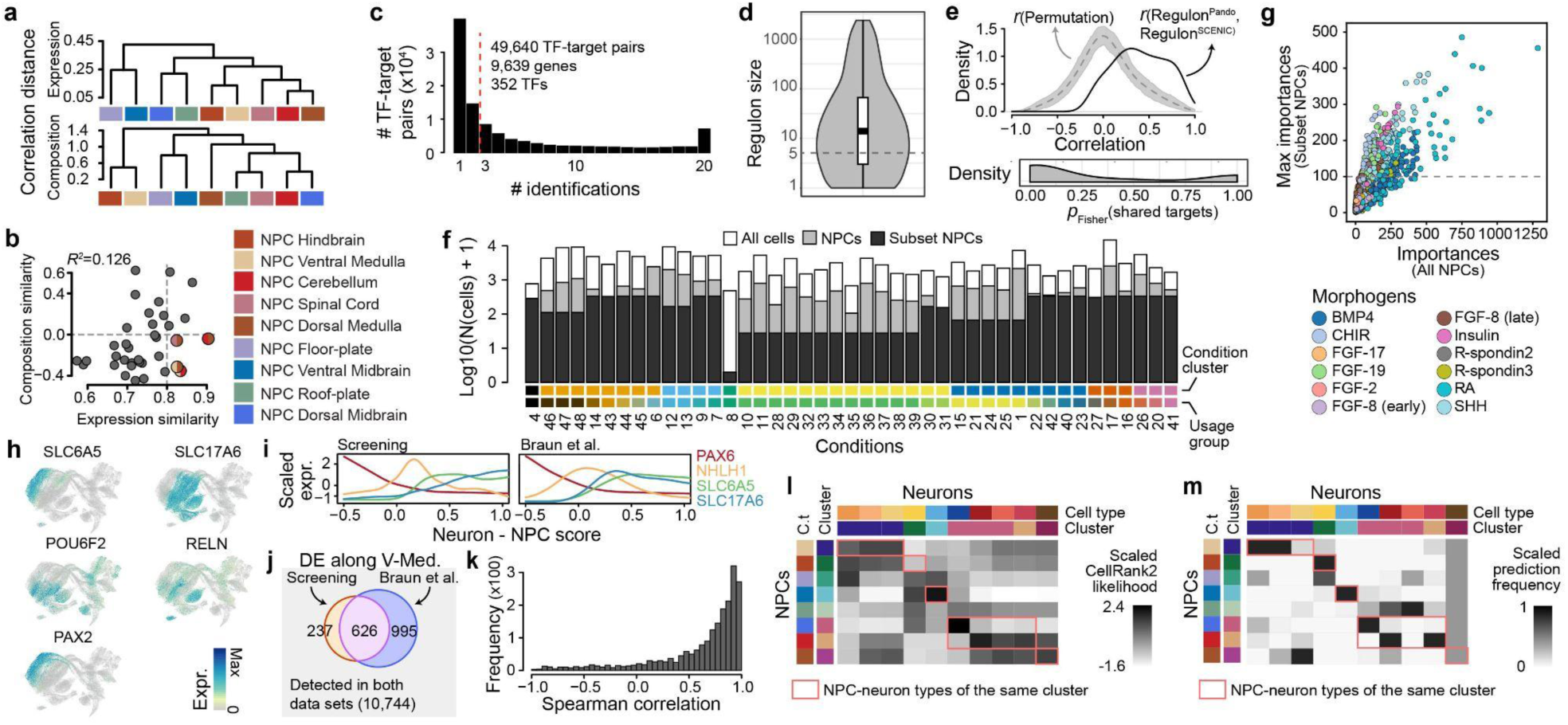
Supplementary analysis of trajectory analysis and inference of regulatory mechanism by morphogens. (a) Dendrograms showing similarities between NPC cell types, based on gene expression distances (upper) or co-occurrence distances (lower). (b) Correlations between expression and co-occurrence similarities. Every dot represents one pair of NPC cell types. The two dashed lines show thresholds to define high and low similarities. Pairs with high expression similarities but low co-occurrence similarities are shown as two semicircles, each of which is colored by a NPC cell type of the pair. (c) Frequency of TF-target pairs being identified among SCENIC results on the 20 different runs with different random subsets of NPCs. (d) Distribution of regulon sizes, i.e. numbers of predicted targets per TF. The dash line shows the size threshold for the final regulon list. (e) Consistency of regulon inference between the SCENIC results and the previous Pando results. The upper panel shows distribution of correlation between regulon activities estimated based on the SCENIC prediction and the previous Pando-based inference. The area in shade shows the distribution of 100 permutations. The lower panel shows the one-sided Fisher’s exact test p-value distribution for overlap of predicted targets per TF with the two approaches. P-value smaller than 0.5 indicates more overlap than expected. (f) Numbers of all cells, all NPCs, and targeted NPCs in each condition for GRNBoost2 to infer morphogen-regulon relationship. (g) Correlation between estimated importances for morphogen-regulon pairs by GRNBoost2, using all NPCs (x-axis) or taking the maximal values across 50 runs on different random subsets of NPCs with the targeted numbers (y-axis). Dots are colored by the morphogen in each pair. (h) UMAP of the matched subset of primary atlas to the screening data, colored by selected markers of the ventral medulla cell types. (i) Expression profiles of selected marker genes across the ventral medulla NPC-neuron differentiation scores, in the screening data (left) and the primary data (right). (j) Numbers of differentially expressed genes (DEGs) along the ventral medulla differentiation in the screening data and the primary atlas, as well as their overlap. (k) Distribution of Spearman correlation between differentiation-dependent expression profiles of the identified DEGs in the two systems. (l-m) Scaled differentiation likelihood of different NPCs towards different neuron cell types in the primary atlas, inferred using CellRank2 (l) or stepwise elastic net prediction along the differentiation trajectories (m).

## Supplementary Tables

1) Overview of single-cell genomic experiments.
2) Differential expression between primary and organoid-derived dopaminergic neurons. P values were derived using ANOVA and FDR correction.
3) Peak characterisation.
4) Peak cluster characterisation, including number of regions per cluster and top 10 enriched GO-terms, derived from GREAT analysis and top 10 enriched motifs.
5) Overview of gRNA sequences, primers for gRNA vector generation and pooling strategy for KO experiments.
6) Transcriptomic KO effects in the CRISPR-perturbation screens for *EBF3*-KO, day 30 and *ONCECUT2*-KO, d70. P values were derived using the Wilcoxon test and FDR correction.
7) Experimental conditions for the probing of morphogen combinations in human neural posterior organoids. Dose and treatment windows of each morphogen are stated in separate columns, with the culture days indicated at the top. The number of assayed organoids per morphogen treatment is indicated in the last column.
8) Enrichment effects in the morphogen screen. P values were derived using a t-test and FDR correction.
9) MERSCOPE gene panel.

## References

1. Miles, G. B. & Wyart, C. Editorial overview: Motor control systems of the spinal cord and hindbrain. Curr. Opin. Physiol. 8, iii–v (2019).

2. Doherty, D., Millen, K. J. & Barkovich, A. J. Midbrain and hindbrain malformations: advances in clinical diagnosis, imaging, and genetics. Lancet Neurol 12, 381–393 (2013).

3. Eiraku, M. et al. Self-organized formation of polarized cortical tissues from ESCs and its active manipulation by extrinsic signals. Cell Stem Cell 3, 519–532 (2008).

4. Lancaster, M. A. et al. Cerebral organoids model human brain development and microcephaly. Nature 501, 373–379 (2013).

5. Camp, J. G. et al. Human cerebral organoids recapitulate gene expression programs of fetal neocortex development. Proc Natl Acad Sci U S A 112, 15672–15677 (2015).

6. Klingler, E., Francis, F., Jabaudon, D. & Cappello, S. Mapping the molecular and cellular complexity of cortical malformations. Science 371, (2021).

7. Li, C. et al. Author Correction: Single-cell brain organoid screening identifies developmental defects in autism. Nature 623, E20 (2023).

8. Tieng, V. et al. Engineering of midbrain organoids containing long-lived dopaminergic neurons. Stem cells and development 23, (2014).

9. Kim, H. et al. Modeling G2019S-LRRK2 Sporadic Parkinson’s Disease in 3D Midbrain Organoids. Stem cell reports 12, (2019).

10. Nickels, S. L. et al. Reproducible generation of human midbrain organoids for in vitro modeling of Parkinson’s disease. Stem Cell Res 46, 101870 (2020).

11. Fiorenzano, A. et al. Single-cell transcriptomics captures features of human midbrain development and dopamine neuron diversity in brain organoids. Nature Communications 12, 1–19 (2021).

12. Mohamed, N. et al. Generation of human midbrain organoids from induced pluripotent stem cells. MNI Open Res (2021) doi:10.12688/mniopenres.12816.2.

13. Qian, X. et al. Brain-Region-Specific Organoids Using Mini-bioreactors for Modeling ZIKV Exposure. Cell 165, 1238–1254 (2016).

14. Jo, J. et al. Midbrain-like Organoids from Human Pluripotent Stem Cells Contain Functional Dopaminergic and Neuromelanin-Producing Neurons. Cell Stem Cell 19, 248–257 (2016).

15. Valiulahi, P. et al. Generation of caudal-type serotonin neurons and hindbrain-fate organoids from hPSCs. Stem Cell Reports 16, 1938 (2021).

16. Andersen, J. et al. Generation of Functional Human 3D Cortico-Motor Assembloids. Cell 183, (2020).

17. Muguruma, K., Nishiyama, A., Kawakami, H., Hashimoto, K. & Sasai, Y. Self-organization of polarized cerebellar tissue in 3D culture of human pluripotent stem cells. Cell reports 10, (2015).

18. Atamian, A. et al. Human cerebellar organoids with functional Purkinje cells. Cell Stem Cell 31, 39–51.e6 (2024).

19. Eura, N. et al. Brainstem Organoids From Human Pluripotent Stem Cells. Front. Neurosci. 14, 514037 (2020).

20. Lui, K. N.-C., Li, Z., Lai, F. P.-L., Lau, S.-T. & Ngan, E. S.-W. Organoid models of breathing disorders reveal patterning defect of hindbrain neurons caused by PHOX2B-PARMs. Stem Cell Reports 18, 1500–1515 (2023).

21. He, Z. et al. An integrated transcriptomic cell atlas of human neural organoids. Nature 635, 690–698 (2024).

22. Scuderi, S. et al. Specification of human regional brain lineages using orthogonal gradients of WNT and SHH in organoids. bioRxiv 2024.05.18.594828 (2024) doi:10.1101/2024.05.18.594828.

23. Sanchís-Calleja, F. et al. Decoding morphogen patterning of human neural organoids with a multiplexed single-cell transcriptomic screen. bioRxiv 2024.02.08.579413 (2024) doi:10.1101/2024.02.08.579413.

24. Amin, N. D. et al. Generating human neural diversity with a multiplexed morphogen screen in organoids. Cell Stem Cell 31, 1831–1846.e9 (2024).

25. Qian, X. et al. Generation of human brain region-specific organoids using a miniaturized spinning bioreactor. Nat Protoc 13, 565–580 (2018).

26. Braun, E. et al. Comprehensive cell atlas of the first-trimester developing human brain. Science 382, eadf1226 (2023).

27. Lotfollahi, M. et al. Mapping single-cell data to reference atlases by transfer learning. Nat Biotechnol 40, 121–130 (2022).

28. Bravo González-Blas, C., et al. cisTopic: cis-regulatory topic modeling on single-cell ATAC-seq data. Nat Methods 16, 397–400 (2019).

29. De Donno, C. et al. Population-level integration of single-cell datasets enables multi-scale analysis across samples. Nat Methods 20, 1683–1692 (2023).

30. Mannens, C. C. A. et al. Chromatin accessibility during human first-trimester neurodevelopment. Nature (2024) doi:10.1038/s41586-024-07234-1.

31. Fleck, J. S. et al. Inferring and perturbing cell fate regulomes in human brain organoids. Nature 621, 365–372 (2023).

32. Vernay, B. et al. Otx2 regulates subtype specification and neurogenesis in the midbrain. J Neurosci 25, 4856–4867 (2005).

33. Dang, N. D. P. et al. Disrupted development of sensory systems and the cerebellum in a zebrafish ebf3a mutant. Neuroscience (2024).

34. Mall, M. et al. Myt1l safeguards neuronal identity by actively repressing many non-neuronal fates. Nature 544, 245–249 (2017).

35. Chen, J., Fuhler, N. A., Noguchi, K. K. & Dougherty, J. D. MYT1L is required for suppressing earlier neuronal development programs in the adult mouse brain. Genome Res 33, 541–556 (2023).

36. van der Raadt, J., van Gestel, S. H. C., Nadif Kasri, N. & Albers, C. A. ONECUT transcription factors induce neuronal characteristics and remodel chromatin accessibility. Nucleic Acids Res 47, 5587–5602 (2019).

37. Rosenberg, A. B. et al. Single-cell profiling of the developing mouse brain and spinal cord with split-pool barcoding. Science 360, 176–182 (2018).

38. Jordan, J., Böttner, M., Schluesener, H. J., Unsicker, K. & Krieglstein, K. Bone morphogenetic proteins: neurotrophic roles for midbrain dopaminergic neurons and implications of astroglial cells. Eur J Neurosci 9, 1699–1709 (1997).

39. Induction of a dopaminergic phenotype in cultured striatal neurons by bone morphogenetic proteins. Developmental Brain Research 130, 91–98 (2001).

40. Lin, H.-C. et al. Human neuron subtype programming through combinatorial patterning with scRNA-seq readouts. bioRxiv 2023.12.12.571318 (2023) doi:10.1101/2023.12.12.571318.

41. Aibar, S. et al. SCENIC: single-cell regulatory network inference and clustering. Nat Methods 14, 1083–1086 (2017).

42. Moerman, T. et al. GRNBoost2 and Arboreto: efficient and scalable inference of gene regulatory networks. Bioinformatics 35, 2159–2161 (2019).

43. Weiler, P., Lange, M., Klein, M., Pe’er, D. & Theis, F. CellRank 2: unified fate mapping in multiview single-cell data. Nat. Methods 21, 1196–1205 (2024).

44. Toh, H. S. Y. et al. BrainSTEM: A multi-resolution fetal brain atlas to assess the fidelity of human midbrain cultures. bioRxiv (2024) doi:10.1101/2024.09.17.613274.

45. Nolbrant, S., et al. Interspecies organoids reveal human-specific molecular features of dopaminergic neuron development and vulnerability. bioRxivorg (2024) doi:10.1101/2024.11.14.623592.

46. Ge, M. et al. A Spacetime Odyssey of Neural Progenitors to Generate Neuronal Diversity. Neurosci Bull 39, 645–658 (2023).

47. Missarova, A. et al. geneBasis: an iterative approach for unsupervised selection of targeted gene panels from scRNA-seq. Genome Biol 22, 333 (2021).

48. Kanton, S. et al. Organoid single-cell genomic atlas uncovers human-specific features of brain development. Nature 574, 418–422 (2019).

49. He, Z. et al. Lineage recording in human cerebral organoids. Nat Methods 19, 90–99 (2022).

50. Jain, A. et al. Morphodynamics of human early brain organoid development. bioRxiv 2023.08.21.553827 (2023) doi:10.1101/2023.08.21.553827.

51. He, Z., Brazovskaja, A., Ebert, S., Camp, J. G. & Treutlein, B. CSS: cluster similarity spectrum integration of single-cell genomics data. Genome Biol 21, 224 (2020).

52. González, F. et al. An iCRISPR platform for rapid, multiplexable, and inducible genome editing in human pluripotent stem cells. Cell Stem Cell 15, 215–226 (2014).

53. Zhu, Z., González, F. & Huangfu, D. The iCRISPR platform for rapid genome editing in human pluripotent stem cells. Methods Enzymol. 546, 215–250 (2014).

54. Stuart, T. et al. Comprehensive Integration of Single-Cell Data. Cell 177, 1888–1902.e21 (2019).

55. Kang, H. M. et al. Author Correction: Multiplexed droplet single-cell RNA-sequencing using natural genetic variation. Nat Biotechnol 38, 1356 (2020).

56. McInnes, L., Healy, J., Saul, N. & Großberger, L. UMAP: Uniform Manifold Approximation and Projection. J. Open Source Softw. 3, 861 (2018).

57. Garritsen, O., van Battum, E. Y., Grossouw, L. M. & Pasterkamp, R. J. Development, wiring and function of dopamine neuron subtypes. Nat Rev Neurosci 24, 134–152 (2023).

58. La Manno, G. et al. Molecular Diversity of Midbrain Development in Mouse, Human, and Stem Cells. Cell 167, 566–580.e19 (2016).

59. Yaghmaeian Salmani, B., et al. Transcriptomic atlas of midbrain dopamine neurons uncovers differential vulnerability in a Parkinsonism lesion model. (2024) doi:10.7554/elife.89482.2.

60. Yang, K. et al. CHRNA5 gene variation affects the response of VTA dopaminergic neurons during chronic nicotine exposure and withdrawal. Neuropharmacology 235, 109547 (2023).

61. Poulin, J.-F., Gaertner, Z., Moreno-Ramos, O. A. & Awatramani, R. Classification of Midbrain Dopamine Neurons Using Single-Cell Gene Expression Profiling Approaches. Trends Neurosci 43, 155–169 (2020).

62. Pachitariu, M. & Stringer, C. Cellpose 2.0: how to train your own model. Nat Methods 19, 1634–1641 (2022).

63. Gu, Z. & Hübschmann, D. rGREAT: an R/bioconductor package for functional enrichment on genomic regions. Bioinformatics 39, (2023).

64. Heinz, S. et al. Simple combinations of lineage-determining transcription factors prime cis-regulatory elements required for macrophage and B cell identities. Mol Cell 38, 576–589 (2010).

65. Rauluseviciute, I. et al. JASPAR 2024: 20th anniversary of the open-access database of transcription factor binding profiles. Nucleic Acids Res 52, D174–D182 (2024).

66. Riddell, A., Hopper, T., Shaoze, L. U. O., Leinweber, K. & Grivas, A. Lda-Project/lda: 1.1.0. (Zenodo, 2018). doi:10.5281/ZENODO.1412135.

67. Schep, A. N., Wu, B., Buenrostro, J. D. & Greenleaf, W. J. chromVAR: inferring transcription-factor-associated accessibility from single-cell epigenomic data. Nat Methods 14, 975–978 (2017).

68. Papalexi, E. et al. Characterizing the molecular regulation of inhibitory immune checkpoints with multimodal single-cell screens. Nat. Genet. 53, 322–331 (2021).

69. Alquicira-Hernandez, J. & Powell, J. E. Nebulosa recovers single-cell gene expression signals by kernel density estimation. Bioinformatics 37, 2485–2487 (2021).

70. Yu, G., Wang, L.-G., Han, Y. & He, Q.-Y. clusterProfiler: an R package for comparing biological themes among gene clusters. OMICS 16, 284–287 (2012).

71. Wolf, F. A., Angerer, P. & Theis, F. J. SCANPY: large-scale single-cell gene expression data analysis. Genome Biol 19, 15 (2018).

72. Gray, P. A. Transcription factors define the neuroanatomical organization of the medullary reticular formation. Front Neuroanat 7, 7 (2013).

73. Lahti, L., Achim, K. & Partanen, J. Molecular regulation of GABAergic neuron differentiation and diversity in the developing midbrain. Acta Physiol (Oxf) 207, 616–627 (2013).

74. Hoshino, M. Neuronal subtype specification in the cerebellum and dorsal hindbrain. Dev Growth Differ 54, 317–326 (2012).

75. Hernandez-Miranda, L. R., Müller, T. & Birchmeier, C. The dorsal spinal cord and hindbrain: From developmental mechanisms to functional circuits. Dev Biol 432, 34–42 (2017).

76. Bertuzzi, S. et al. Characterization of Lhx9, a novel LIM/homeobox gene expressed by the pioneer neurons in the mouse cerebral cortex. Mech Dev 81, 193–198 (1999).

77. Kozareva, V. et al. A transcriptomic atlas of mouse cerebellar cortex comprehensively defines cell types. Nature 598, 214–219 (2021).

78. Carmichael, K., Sullivan, B., Lopez, E., Sun, L. & Cai, H. Diverse midbrain dopaminergic neuron subtypes and implications for complex clinical symptoms of Parkinson’s disease. Ageing Neurodegener Dis 1, (2021).

79. Phipson, B. et al. propeller: testing for differences in cell type proportions in single cell data. Bioinformatics 38, 4720–4726 (2022).

80. Xu, C. et al. Probabilistic harmonization and annotation of single-cell transcriptomics data with deep generative models. Mol Syst Biol 17, e9620 (2021).

81. Lopez, R., Regier, J., Cole, M. B., Jordan, M. I. & Yosef, N. Deep generative modeling for single-cell transcriptomics. Nat Methods 15, 1053–1058 (2018).

82. La Manno, G. et al. Molecular architecture of the developing mouse brain. Nature 596, 92–96 (2021).

83. Korsunsky, I. et al. Fast, sensitive and accurate integration of single-cell data with Harmony. Nat Methods 16, 1289–1296 (2019).

84. Herman, J. S., Sagar & Grün, D. FateID infers cell fate bias in multipotent progenitors from single-cell RNA-seq data. Nat Methods 15, 379–386 (2018).

